# Rare disease research workflow using multilayer networks elucidates the molecular determinants of severity in Congenital Myasthenic Syndromes

**DOI:** 10.1101/2023.01.19.524736

**Authors:** Iker Núñez-Carpintero, Emily O’Connor, Maria Rigau, Mattia Bosio, Sally Spendiff, Yoshiteru Azuma, Ana Topf, Rachel Thompson, Peter A.C. ’t Hoen, Teodora Chamova, Ivailo Tournev, Velina Guergueltcheva, Steven Laurie, Sergi Beltran, Salvador Capella, Davide Cirillo, Hanns Lochmüller, Alfonso Valencia

## Abstract

Exploring the molecular basis of disease severity in rare disease scenarios is a challenging task provided the limitations on data availability. Causative genes have been described for Congenital Myasthenic Syndromes (CMS), a group of diverse minority neuromuscular junction (NMJ) disorders; yet a molecular explanation for the phenotypic severity differences remains unclear. Here, we present a workflow to explore the functional relationships between CMS causal genes and altered genes from each patient, based on multilayer network analysis of protein-protein interactions, pathways and metabolomics.

Our results show that CMS severity can be ascribed to the personalized impairment of extracellular matrix components and postsynaptic modulators of acetylcholine receptor (AChR) clustering. We explore this in more detail for one of the proteins not previously associated with the NMJ, USH2A. Loss of the zebrafish USH2A ortholog revealed some effects on early movement and gross NMJ morphology.

This work showcases how coupling multilayer network analysis with personalized -omics information provides molecular explanations to the varying severity of rare diseases; paving the way for sorting out similar cases in other rare diseases.

## Introduction

Understanding phenotypic severity is crucial for prediction of disease outcomes, as well as for administration of personalized treatments. Different severity levels among patients presenting the same medical condition could be explained by characteristic relationships between diverse molecular entities (i.e. gene products, metabolites, etc) in each individual. In this setting, multi-omics data integration is becoming a promising tool for research, as it has the potential to gain complex insights of the molecular determinants underlying disease heterogeneity. However, even in a scenario where the level of biomedical detail available to study is growing in an exponential manner (Karczewski and Snyder, 2018), the analysis of the molecular determinants of disease severity is not typically addressed in rare disease research literature (Boycott et al., 2013), despite its obvious relevance at the medical and clinical level. Rare diseases represent a challenging setting for the application of precision medicine because, by definition, they affect a small number of patients, and therefore the data available for study is considerably limited in comparison to other conditions. Accordingly, leveraging the wealth of biomedical knowledge of diverse nature coming from publicly available databases has the potential to address data limitations in rare diseases (Buphamalai et al., 2021; Mitani and Haneuse, 2020). In this sense, multilayer networks can offer a holistic representation of biomedical data resources (Gosak et al., 2018; Halu et al., 2019), which may allow exploration of the biology related to a given disease independently of cohort sizes and their available omics data.

Here, in order to evaluate and demonstrate the potential of multilayer networks as means of assessing severity in rare disease scenarios, we provide an illustrative case where we develop a framework for analyzing a patient cohort affected by Congenital Myasthenic Syndromes (CMS), a group of inherited rare disorders of the neuromuscular junction (NMJ). Fatigable weakness is a common hallmark of these syndromes, that affects approximately 1 patient in 150,000 people worldwide. The inheritance of CMS is autosomal recessive in the majority of patients. CMS can be considered a relevant use case because, while patients share similar clinical and genetic features (Finsterer, 2019), phenotypic severity of CMS varies greatly, with patients experiencing a range of muscle weakness and movement impairment. While over 30 genes are known to be monogenic causes of different forms of CMS (**Table 1**), these genes do not fully explain the ample range of observed severities, which has been suggested to be determined by additional factors involved in neuromuscular function (Thompson et al. 2019). Examples of CMS-related genes are *AGRN, LRP4* and *MUSK* which code for proteins that mediate communication between the nerve ending and the muscle, which is crucial for formation and maintenance of the NMJ **(Figure 1)**.

**Table 1.**
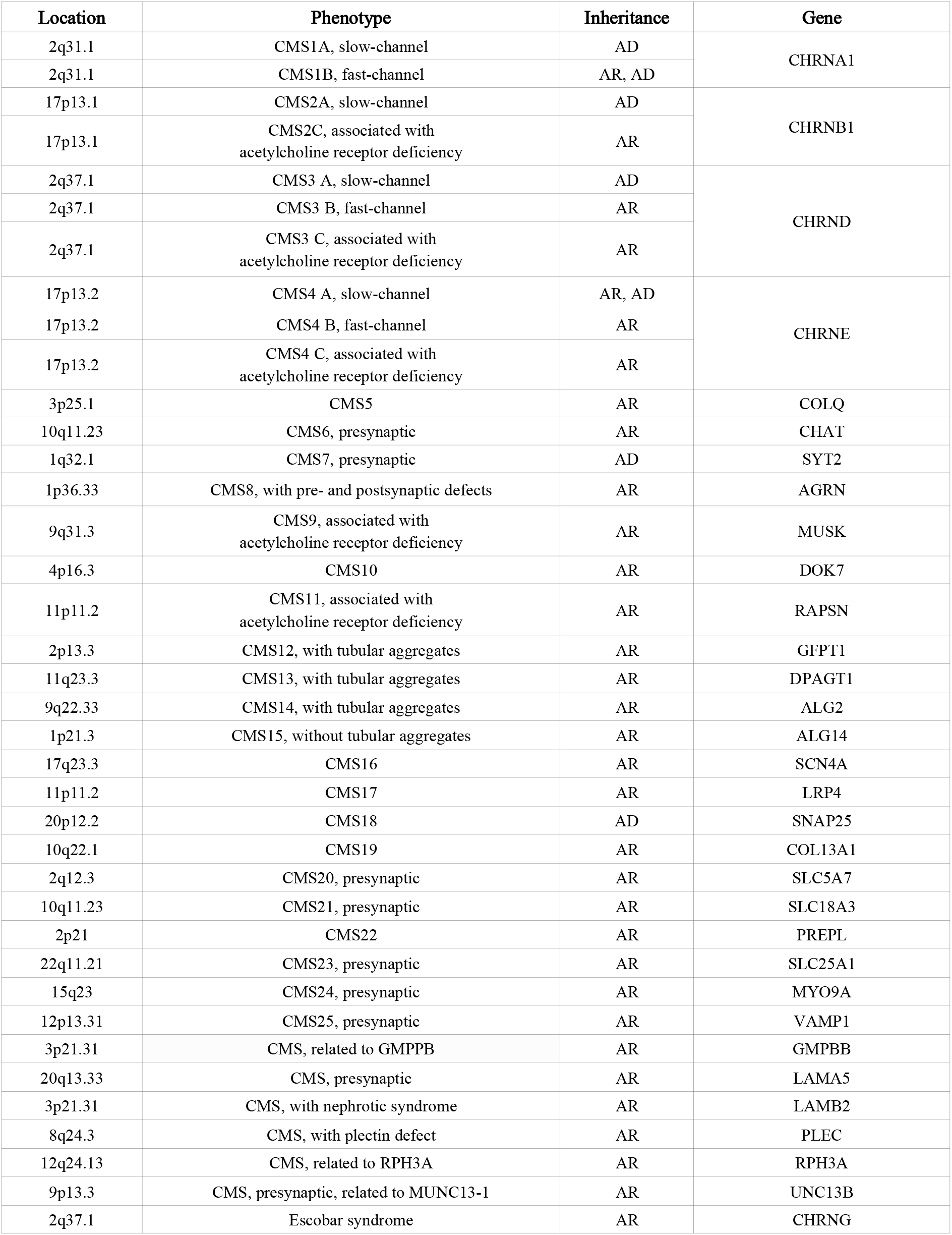
Location, phenotype, inheritance and genes involved in CMS (adapted from https://omim.org/phenotypicSeries/PS601462 and http://www.musclegenetable.fr). AR: autosomal recessive; AD: autosomal dominant.

**Figure 1.**
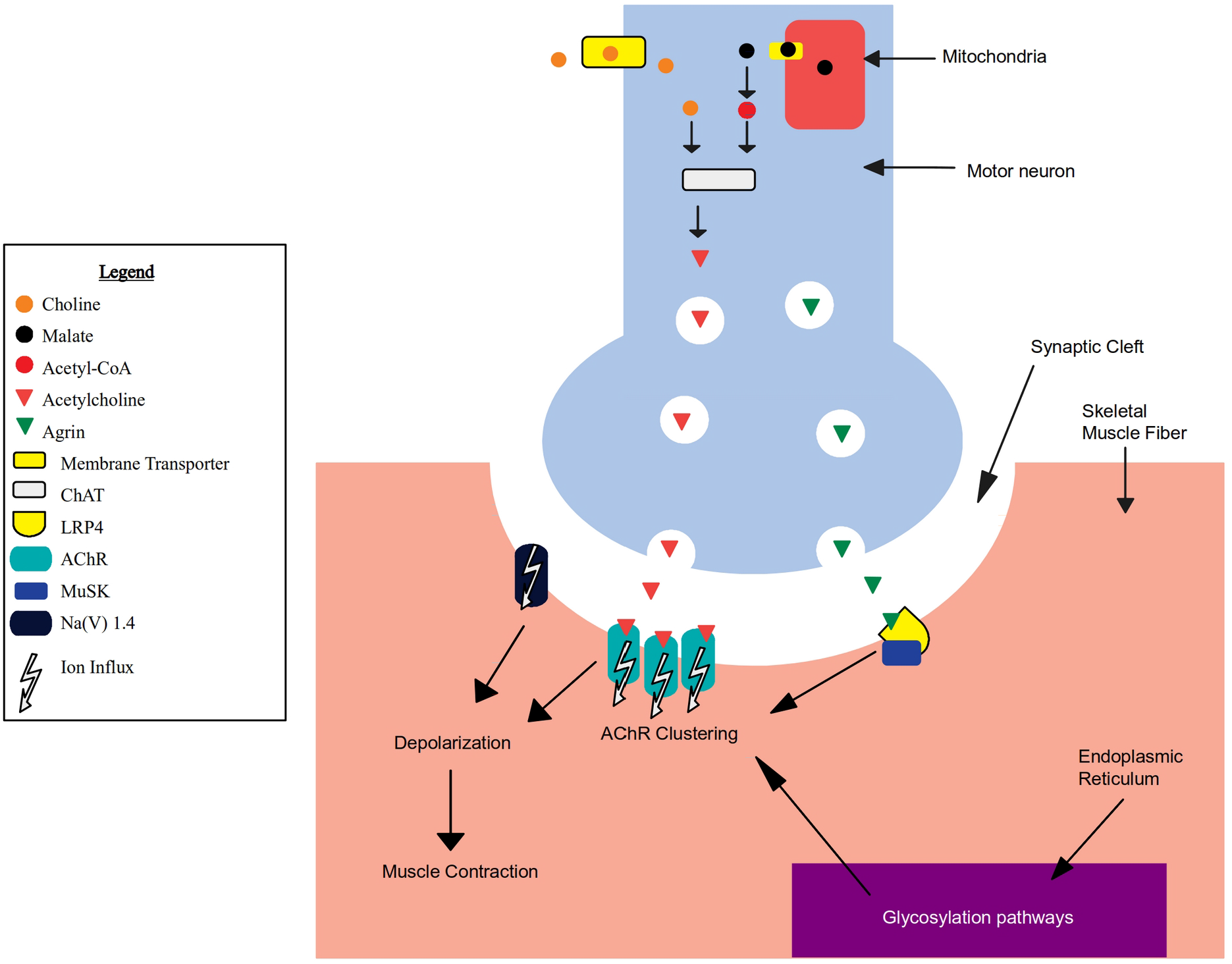
A schematic depiction of the main molecular activities of known CMS causal genes (Methods) taking place at the neuromuscular junction (NMJ) in the presynaptic terminal (in blue), synaptic cleft (in white), and skeletal muscle fiber (in yellow) (for a detailed description of this system see **Supplementary Information**).

In particular, the AGRN-LRP4 receptor complex activates MUSK by phosphorylation, inducing clustering of the acetylcholine receptor (AChR) in the postsynaptic membrane allowing the presynaptic release of acetylcholine (ACh) to trigger muscle contraction (Burden et al., 2013; Li et al., 2018). Additional evidence of CMS severity heterogeneity emerged within the NeurOmics and RD-Connect projects (Lochmüller et al., 2018) studying a small population (about 100 individuals) of gypsy ethnic origin from Bulgaria.

All affected individuals shared the same causal homozygous mutation (a deletion within the AChR ε subunit, *CHRNE* c.1327delG (Abicht et al., 1999)), however, the severity of symptoms across this cohort varies considerably regardless of age, gender and initiated therapy, suggesting the existence of additional genetic causes for the diversity of disease phenotypes. By analyzing multi-omics data, we performed an in-depth characterization of 20 CMS patients, representing the two opposite ends of the spectrum observed in the wider cohort, aiming to investigate the molecular basis of the observed differences in the individual severity of the disease. Clinically, CMS severity ranges from minor symptoms (e.g., exercise intolerance) to more severe CMS forms and it is dependent on the causal genetic impairments (Abicht et al., 1993; Della Marina et al., 2020). Severe CMS is typically presented with reduced Forced Vital Capacity (FVC), severe generalized muscle fatigue and weakness, proximal and bulbar muscle fatigue and weakness, impaired myopathic gait and hyperlordosis. Two CMS severity levels have been identified for this cohort through extensive phenotyping, namely a severe disease phenotype (8 patients) and a not-severe disease phenotype (2 intermediate and 10 mild patients) (**Suppl. Table 1**). Out of the tested demographic factors (age, sex) and clinical tests (speech, mobility, respiratory dysfunctions, among others), FVC and shoulder lifting ability show a significant association with the severity classes (two-tailed Fisher’s exact test p-values of 0.0128 and 0.0418, respectively; **Suppl. Figure 1**). We sought to interrogate whether severity was determined by additional genetic variations impacting neuromuscular activity, on top of the causative CHRNE mutation. We analyzed three main types of genetic variations: single nucleotide polymorphisms (SNPs), copy number variations (CNVs), and compound heterozygous variants (two recessive alleles located at different loci within the same gene in a given individual). The extensive analysis of the genomic information did not render any SNPs that could be considered a unique cause of disease severity by being common to all the cases. Nevertheless, a number of CNVs and compound heterozygous variants were found to appear exclusively in the different severity groups, in one or more patients. Moreover, the compound heterozygous variants of the severe group are enriched in pathways related to the extracellular matrix (ECM) receptors, which have been proposed as a target for CMS therapy (Ito and Ohno, 2018).

To investigate the functional relationship between these variants and CMS severity, **we designed an analytical workflow based on multilayer networks (Figure 2)**, allowing the integration of external biological knowledge to acquire deeper functional insights. A multilayer network consists of several layers of nodes and edges describing different aspects of a system (Kivelä et al., 2014). In biomedicine, this data representation has been used to study biomolecular interactions (Zitnik and Leskovec, 2017) and diseases (Halu et al., 2019), facilitating integration and interpretation of heterogeneous sources of data. Several established tools for network analysis have been recently adapted for multilayer networks, such as random walk with restart (Edler et al., 2017; Valdeolivas et al., 2019), community detection algorithms (Didier et al., 2015) and node embeddings (Pio-Lopez et al., 2021). By crossing patient genomic data with the information provided by a biomedical knowledge multilayer network, we are able to describe the functional relationships of new genetic modifiers responsible for the different phenotypic severity levels, showcasing the potential of multilayer networks to provide support on the analysis of rare disease patients.

**Figure 2.**
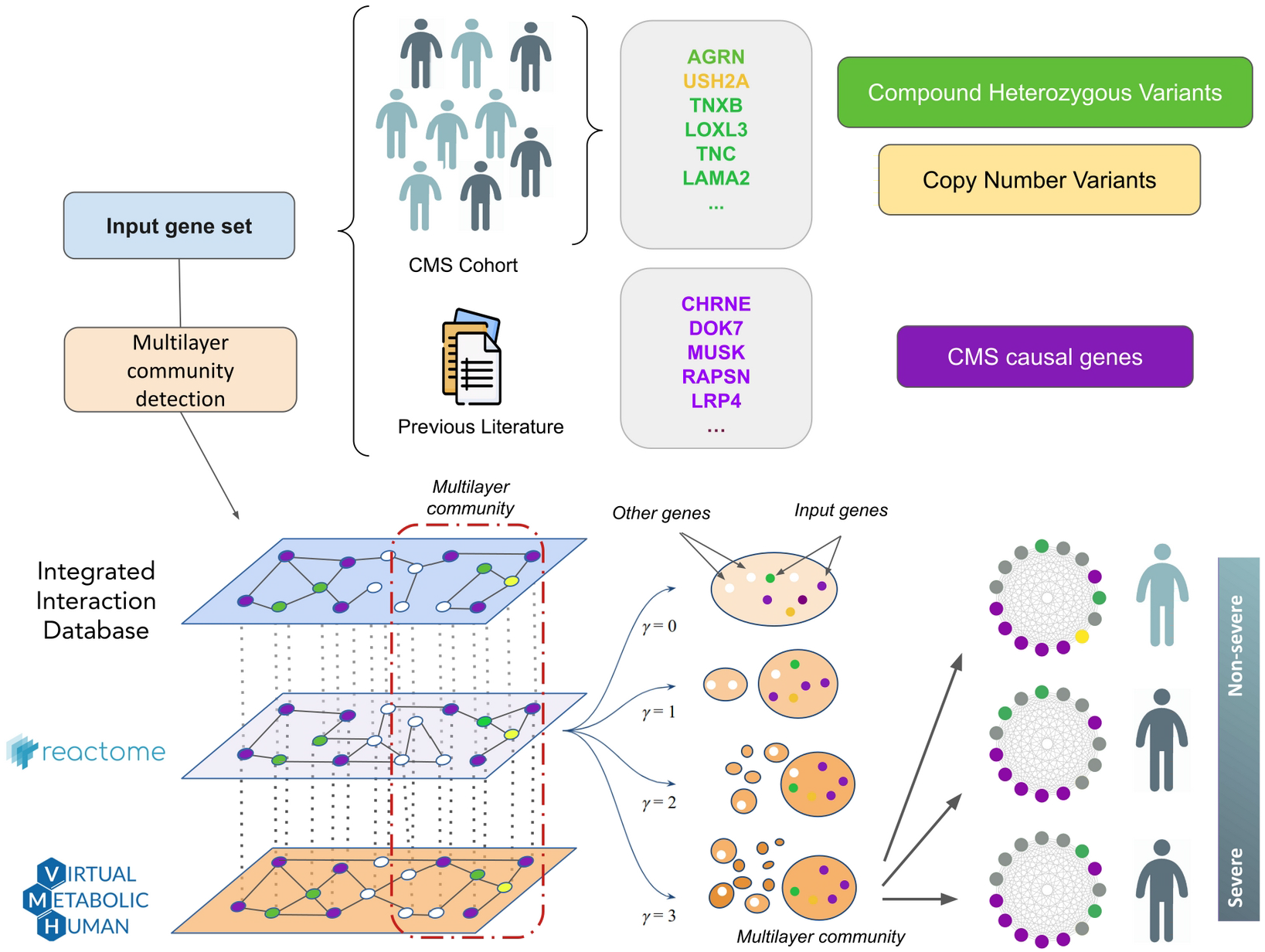
Analytical workflow employed to address the severity of a cohort of patients affected by Congenital Myasthenic Syndromes (CMS). A multi-scale functional analysis approach, based on multilayer networks, was used to identify the functional relationships between genetic alterations obtained from omics data (Whole Genome Sequencing, WGS; RNA-sequencing, RNAseq) with known CMS causal genes. Modules of CMS linked genes are detected using graph community detection at a resolution range (γ) (Methods) where the most prominent changes in community structure occur. Modules that emerged from this analysis were characterized at single individual level.

## Results

### Variants do not segregate with patient severity

We first searched for variants able to segregate the disease phenotypes (severe and not-severe) by analyzing a large panel of mutational events (mutations in isoforms, splicing sites, small and long noncoding genes, promoters, TSS, predicted pathogenic mutations, loss of function mutations, among others). We could not find one single mutation or combinations of mutations that were able to completely segregate the two groups (**Supplementary Information**) although partial segregation can be observed (**Suppl. Table 2**). As already described for monogenic diseases (Kousi and Katsanis, 2015) and cancer (Castro-Giner et al., 2015), we hypothesized that distinct weak disease-promoting effects may represent patient-specific causes to CMS severity, which bring damage to sets of genes that are functionally related. To find these causes, we sought to search for variants with the potential to alter gene functions, such as CNVs and compound heterozygous variants, which have been previously reported to be key to CMS (Abicht et al., 1993; Bevilacqua et al., 2017; Richard et al., 2003; Yang et al., 2018).

### Compound heterozygous variants are functionally related

In order to explore the hypothesis that disease severity in this cohort is due to variants in patient-specific critical elements, we sought to identify potentially damaging compound heterozygous variants and CNVs. We analyzed the gene lists associated with these mutations to search for evidence of alterations in relevant pathways for the severe (n=8) and not-severe cases (n=12). We first performed a functional enrichment analysis (Methods) of the genes with CNVs found in the two groups. The set of affected genes in the severe group is composed of 26 unique genes (10 private to the severe group), while the not-severe group presented 86 unique genes (**Suppl. Table 3**). None of these gene sets showed any functional enrichment. Moreover, none of these genes had been described as causal for CMS, and none carried compound heterozygous variants. (**Suppl. Figure 2**). As for compound heterozygous variants, the set of affected genes in the severe group is composed of 112 unique genes (89 private to the severe group), while the not-severe group resulted in 152 unique genes (**Suppl. Table 3**). We found that the severe group shows significant enrichment in genes belonging to extracellular matrix (ECM) pathways, in particular “ECM receptor interactions” (KEGG hsa04512, adjusted p-value 0.002337) and “ECM proteoglycans” (Reactome R-HSA-30001787, adjusted p-value 0.001237), which are the top-hit pathways when the 89 genes appearing only in the severe group are considered. Both these pathways share common genes, namely *TNXB*, *LAMA2*, *TNC*, and *AGRN*. The role of extracellular matrix proteins for the formation and maintenance of the NMJ has recently drawn attention to the study of CMS (Beeson, 2016; Rodríguez Cruz et al., 2018). In particular, within the genes linked with ECM pathways, *AGRN* and *LAMA2* stand out for their implication in CMS and other rare neuromuscular diseases (Bertini et al., 2011; Bönnemann et al., 2014; Nicole et al., 2014). ECM-related pathways are not enriched in the not-severe set of genes (KEGG hsa04512, adjusted p-value 0.6170). Moreover, top-hit pathways of the not-severe set of genes are not explicitly related to ECM and not consistent between Reactome and KEGG (Reactome “Susceptibility to colorectal cancer” R-HSA-5083636, adjusted p-value 4.131e-7, genes *MUC3A/5B/12/16/17/19*; KEGG “Huntington’s disease” hsa05016, adjusted p-value 0.07103, genes *REST, CREB3L4, CLTCL1, DNAH2/8/10/11*). These findings support our hypothesis that the severe patients might present disruptions in NMJ functionally related genes that, combined with *CHRNE* causative alteration, may be responsible for the worsening of symptoms.

### CMS-specific monolayer and multilayer community detection

As disease-related genes tend to be interconnected (Menche et al., 2015), we sought to analyze the relationships among the CMS linked genes (i.e. known CMS causal genes, and severe and not-severe compound heterozygous variants and CNVs; Methods) using network community clustering analysis. We employed the Louvain algorithm (Methods) to find groups of interrelated genes in three monolayer networks that represents biological knowledge contained in databases, separately: the Reactome database (Fabregat et al., 2018), the Recon3D Virtual Metabolic Human database (Brunk et al., 2018) (both downloaded in May 2018), and from the Integrated Interaction Database (IID) (Kotlyar et al., 2019) (downloaded in October 2018) (**Suppl. Figure 3**). The first network consists of 10,618 nodes (genes) and 875,436 edges, representing shared pathways between genes. The second network consists of 1,863 nodes (genes) and 902,188 edges, representing shared reaction metabolites between genes. The third network consists of 18,018 nodes (genes) and 947,606 edges, representing aggregated protein-protein interactions from all tissues (**Methods:** Monolayer community detection). The last two networks, represent the ‘metabolome’ and the ‘interactome’ data, respectively. Measurement of network overlap and community similarity (**Methods**), revealed high specificity of their edges, as well as that the same CMS linked genes did not form the same communities across the different networks (**Suppl. Figure 4**).

These results show that, although disease-related genes are prone to form well-defined communities in distinct networks (Cantini et al., 2015; Goh et al., 2007), different facets of biological information reflect diverse participation modalities of such genes into communities. In order to deliver an integrated analysis of such heterogeneous information, we further consider them as a multilayer network (Gosak et al., 2018) (**Methods**: Monolayer community detection and Multilayer community detection).

### Large-scale multilayer community detection of disease associated genes

We first sought to test the hypothesis that disease-related genes tend to be part of the same communities also in a multilayer network setting. We used the curated gene-disease associations database DisGeNET (Piñero et al., 2017), showing that disease-associated genes are significantly found to be members of the same multilayer communities (Wilcoxon test p-value < 0.001 in a range of resolution parameters described in the Methods). We pre-processed DisGeNET database by filtering out diseases and disease groups with only one associated gene (6,352 diseases), and those whose number of associated genes was more than 1.5 * interquartile range (IQR) of the gene associated per disease distribution (823 diseases with more than 33 associated genes) (**Suppl. Figure 5A-B**).

This procedure prevents a possible analytical bias due to the higher amounts of genes annotated to specific disease groups (e.g. entry C4020899, “Autosomal recessive predisposition”, annotates 1445 genes). We then retrieved the communities of each associated gene, excluding 428 genes not present in our multilayer network and the diseases left with only one associated gene. The final analysis comprised a total of 5,892 diseases with an average number of 7.38 genes per disease.

For each disease, we counted the number of times that disease-associated genes are found in the same multilayer communities, and compared the distribution of such frequencies with that of balanced random associations (1000 randomizations). Results show that disease-associated genes are significantly found in the same multilayer communities across the resolution interval (**Suppl. Figure 5C**).

### Modules within the CMS multilayer communities

We define a module as a group of CMS linked genes that are systematically found to be part of the same multilayer community while increasing the multilayer network community resolution parameter (Methods; Supplementary Information; **Figures 3-4**).

**Figure 3.**
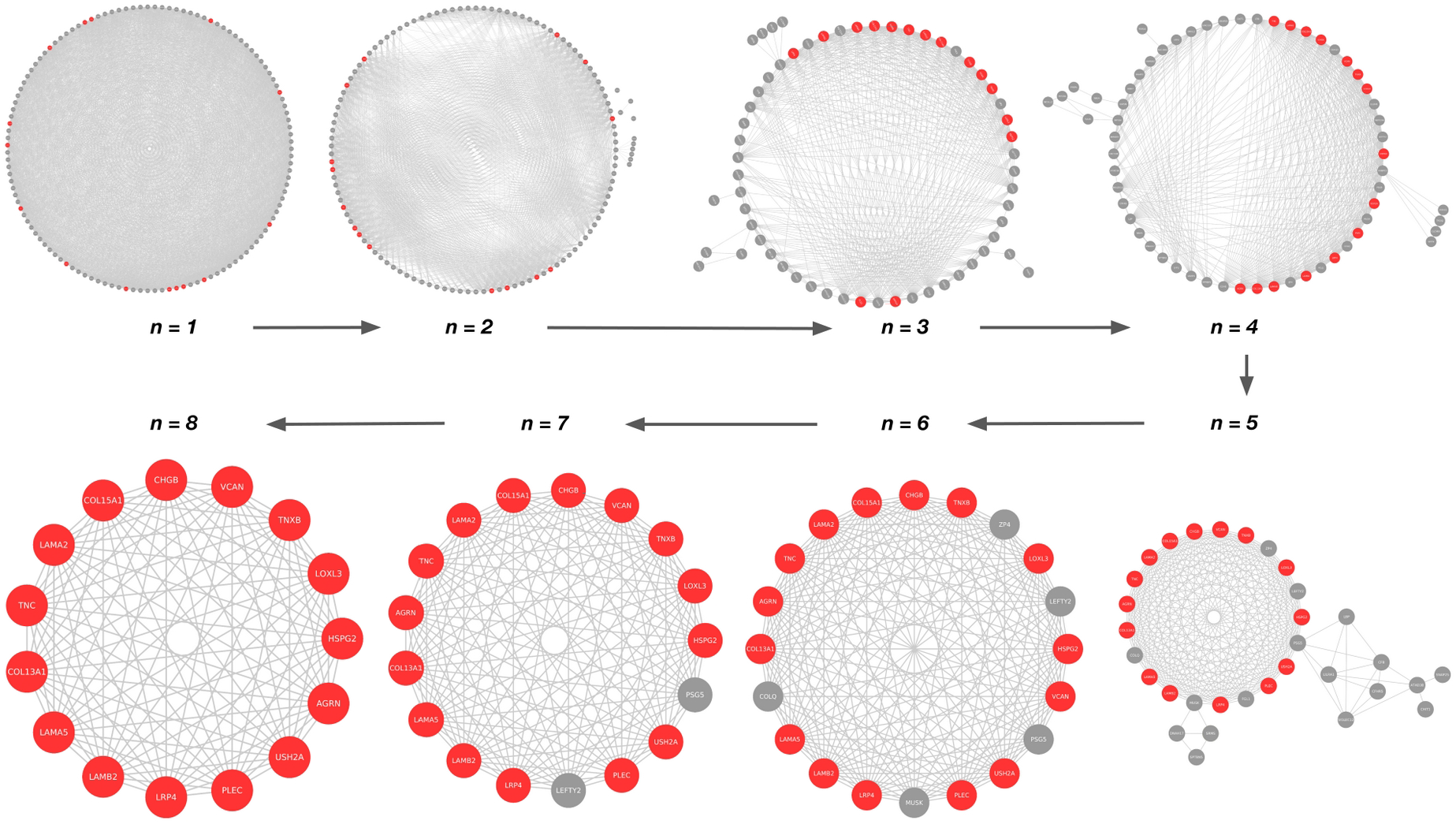
Identification of the largest module containing genes that are found in the same community in the entire range of resolution parameters (Methods). In each module, genes are connected if they are found in the same multilayer communities at *n* values of the resolution parameter γ within the range under consideration (γ∈(0,4]). The arrows indicate the systematic increase of *n*. At *n* = 8, the module contains genes that are always found in the same community in the entire range of resolution (see Supplementary Information "Multilayer community detection analysis"). The largest module containing the CMS linked gene set (highlighted in red), which includes known CMS causal genes, severe-specific heterozygous compound variants and CNVs, is shown.

**Figure 4.**
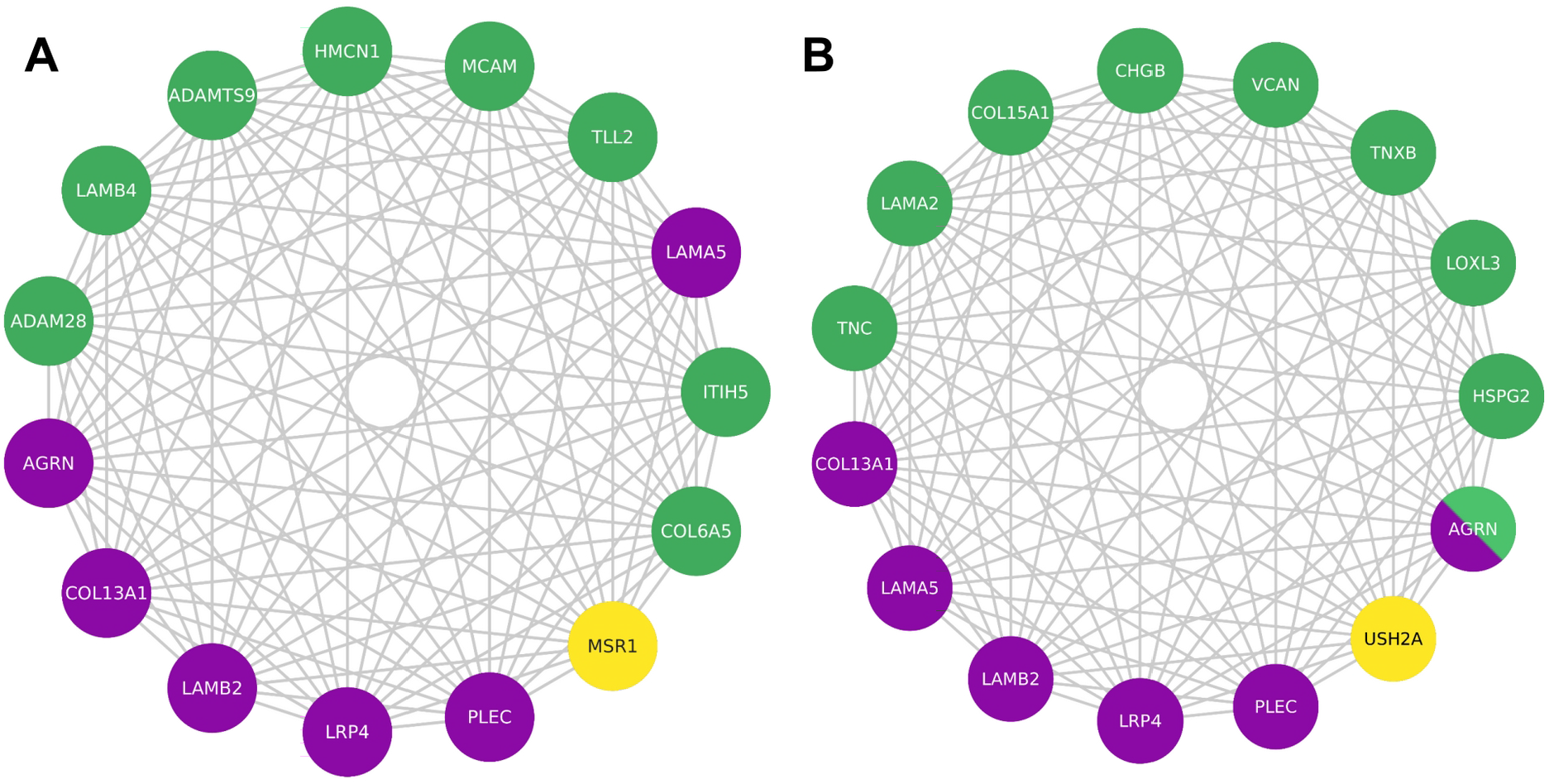
Largest module, containing known CMS causal genes, within the multilayer communities of CMS linked genes that are specific to the not-severe (A) and severe (B) groups. In green, compound heterozygous variants; in yellow, CNVs; in purple, known CMS causal genes. Being a CMS causal gene bearing compound heterozygous variants, AGRN is depicted using both green and purple.

Within each of these communities, we identified smaller modules of CMS linked genes that are specific to the severe and not-severe groups. We tested the significance of obtaining these exact genes in the severe and not-severe largest modules upon severity class label shuffling among all individuals (1000 randomizations). We found that 13 (p-value = 0.022) and 14 (p-value = 0.027) are the minimum number of genes composing the modules that are not expected to be found at random in the severe and not-severe largest components, respectively (**Suppl. Figure 6)**.

In the two groups, the significantly largest module that contains known CMS causal genes is composed of 15 genes (**Figure 4**). 6 out of these 15 are previously described CMS causal genes (Methods), namely the ECM heparan sulfate proteoglycan agrin (*AGRN*); the cytoskeleton component plectin (*PLEC*), causative of myasthenic disease (Forrest et al., 2010); the agrin receptor *LRP4*, key for AChR clustering at NMJ (Barik et al., 2014) and causative of CMS by compound heterozygous variants (Ohkawara et al., 2014); the ECM components *LAMA5* and *LAMB2* laminins, and *COL13A1* collagen. Considering all nodes (not only CMS linked), the number of nodes in the module is 482.

All the other genes of the two modules are involved in a varied spectrum of muscular dysfunctions, discussed in the following sections. As the location of the causal gene products determine the most common classification of the disease (i.e. presynaptic, synaptic, and postsynaptic CMS) (Rodríguez Cruz et al., 2018), we determined class and localization of the members of the found modules (**Table 2**). Laminins, well-known CMS glycoproteins, are affected in both severe (*LAMA2*, *USH2A*) and not-severe (*LAMB4*) groups, and are bound by specific receptors that are damaged in the not-severe group (*MCAM*) (Dagur and McCoy, 2015). Collagens, known CMS-related factors, are associated with the not-severe group (*COL6A5*), and bound by specific receptors that are damaged in the not-severe group (*MSR1*) (Di Martino et al., 2023).

**Table 2.**
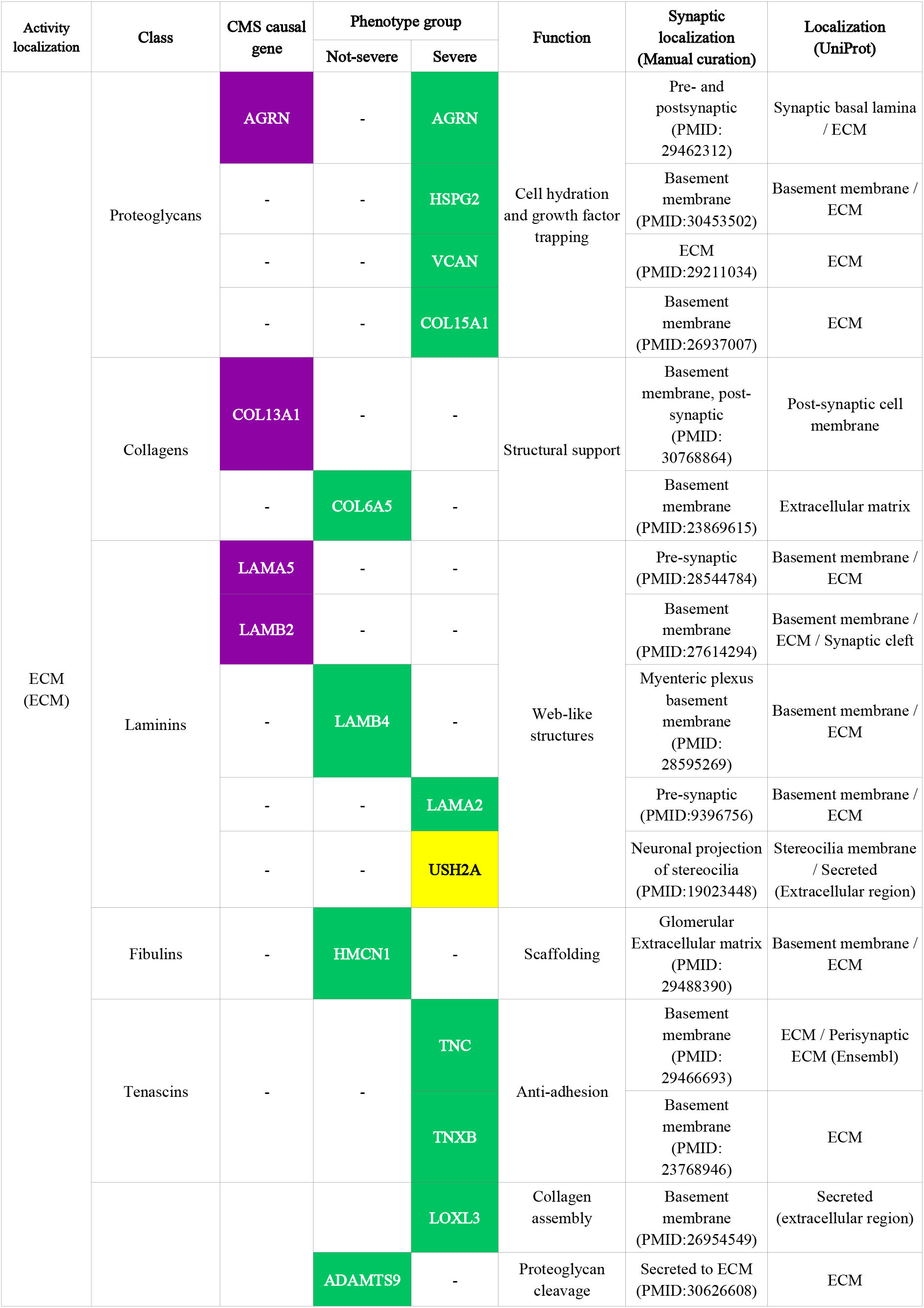
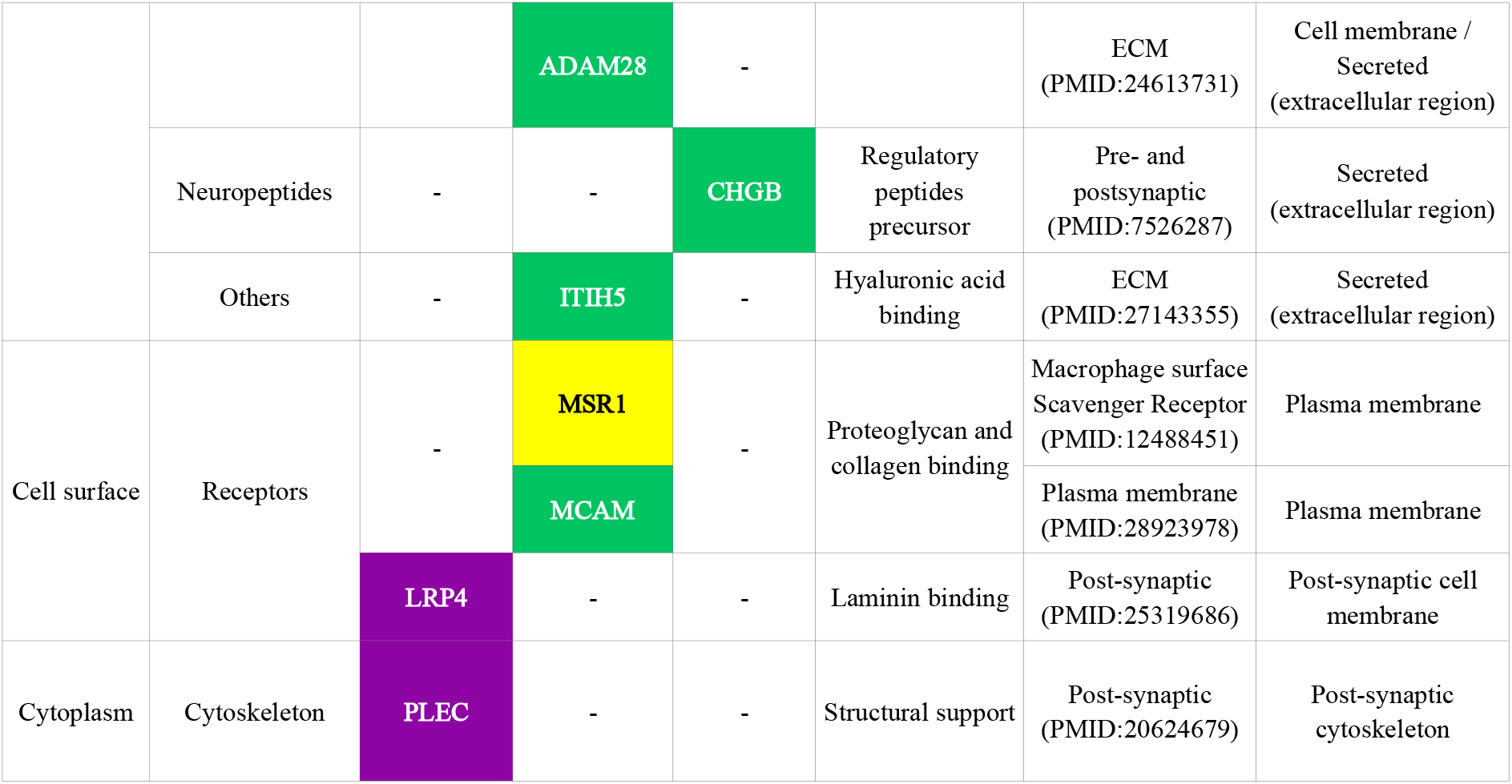
Localization and functions of proteins encoded by the genes found in the largest modules of the multilayer communities of severe and not-severe groups. In green, compound heterozygous variants; in yellow, CNVs; in purple, known CMS causal genes. Synaptic localization was retrieved from manual curation and Uniprot database (Methods).

However, overall collagen biosynthesis is affected in both severe and not-severe groups. Indeed, metalloproteinases, damaged in the not-severe group, are responsible for the proteolytic processing of lysyl oxidases (*LOXL3*), which are implicated in collagen biosynthesis (Panchenko et al., 1996) and damaged in the severe group.

Alterations in proteoglycans (*AGRN*, *HSPG2*, *VCAN*, *COL15A1*) (Iozzo and Schaefer, 2015), tenascins (*TNC*, *TNXB*) (Flück et al., 2008; van Dijk et al., 1993), and chromogranins (*CHGB*) (Andreose et al., 1994) are specific of the severe group. We observed no genes associated with proteoglycan damage in the not-severe group, suggesting a direct involvement of ECM in CMS severity.

### Personalized analysis of the severe cases

We sought to analyze the 15 genes of the largest module of the severe group in each one of the 8 patients, hereafter referred to using the WGS sample labels (**Suppl. Table 1)**. At the topological level, all incident interactions existing between the genes of the severe module (**Figure 4B**) are related to the protein-protein interaction and pathway layers (**Supplementary Figure 7**). Overall, these genes have a varied range of expression levels in tissues of interest (**Suppl. Figure 8**), for instance in skeletal muscle *HSPG2*, *LAMA2*, *PLEC* and *LAMB2* show medium expression levels (9 to 107 TPM) while the others show low expression levels (0.6 to 9 TPM) (Methods).

Patient 2, a 15 years old male, presents compound heterozygous variants in tenascin C (*TNC*), mediating acute ECM response in muscle damage (Flück et al., 2008; Sorensen et al., 2018), and CNVs (specifically, a partial heterozygous copy number loss) in usherin (*USH2A*), which have been associated with hearing and vision loss (Austin-Tse et al., 2018). Patient 16, a 25 years old female, presents compound variants in tenascin XB (*TNXB*), which is mutated in Ehlers-Danlos syndrome, a disease that has already been reported to have phenotypic overlap with muscle weakness (Kirschner et al., 2005; Matsumoto and Aoki, 2020; Okuda-Ashitaka and Matsumoto, 2023; Voermans and Engelen, 2008) and whose compound heterozygous variants have been reported for a primary myopathy case (Pénisson-Besnier et al., 2013; Voermans et al., 2014); and versican (*VCAN*), which has been suggested to modify tenascin C expression (Keller et al., 2012) and is upregulated in Duchenne muscular dystrophy mouse models (McRae et al., 2017, 2020).

Patient 13, a 26 years old male, presents compound mutations in laminin α2 chain (*LAMA2*), a previously reported gene related to various muscle disorders (AMIN et al., 2019; Dimova and Kremensky, 2018; Løkken et al., 2015) whose mutations cause reduction of neuromuscular junction folds (Rogers and Nishimune, 2017), and collagen type XV α chain (*COL15A1*), which is involved in guiding motor axon development (Guillon et al., 2016) and functionally linked to a skeletal muscle myopathy (Eklund et al., 2001; Muona et al., 2002).

Patient 12, a 49 years old female, presents compound mutations in chromogranin B4 (*CHGB*), potentially associated with amyotrophic lateral sclerosis early onset (Gros-Louis et al., 2009; Pampalakis et al., 2019). Patient 18, a 51 years old man, presents compound mutations in agrin (*AGRN*), a CMS causal gene that mediates AChR clustering in the skeletal fiber membrane (Huzé et al., 2009) (Jacquier et al., 2022).

Patient 20, a 57 years old male, presents compound mutations in lysyl oxidase-like 3 (*LOXL3*), involved in myofiber extracellular matrix development by improving integrin signaling through fibronectin oxidation and interaction with laminins (Kraft-Sheleg et al., 2016), and perlecan (*HSPG2*) (Zoeller et al., 2008), a protein present on skeletal muscle basal lamina (Carmen et al., 2019; Larraín et al., 1997), whose deficiency leads to muscular hypertrophy (Xu et al., 2010), that is also mutated in Schwartz-Jampel syndrome (Stum et al., 2006), Dyssegmental dysplasia Silverman-Handmaker type (*DDSH*) (Arikawa-Hirasawa et al., 2001) and fibrosis (Lord et al., 2018), such as Patient 19, a 62 years old female. Furthermore, based on the estimated familial relatedness (Methods) and personal communication (February 2018, Teodora Chamova), patients 19 and 20 are siblings (**Suppl. Table 4**).

### Functional consequences of variants in the severe-specific module

Studying the functional impact of the compound heterozygous variants in the severe-specific genes of the module, we observed that **in 6 of the 8 patients at least one of the variants is predicted to be deleterious by the Ensembl Variant Effect Predictor (VEP)** (McLaren et al., 2016) (Methods; **Suppl. Table 5**). For example, as for Patient 18, who presents 3 different variants in AGRN gene, only rs200607541 is predicted to be deleterious by VEP’s Condel (score = 0.756), SIFT (score = 0.02), and PolyPhen (score = 0.925). In particular, the variant (a C>T transition) presents an allele frequency (AF) of 4.56E-03 (gnomAD exomes) (Karczewski et al., 2020) and affects a region encoding a position related to a EGF-like domain (SMART:SM00181) and a Follistatin-N-terminal like domain (SMART:SM00274). Both of these domains are part of the Kazal domain superfamily which are specially found in the extracellular part of agrins (PFAM: CL0005) (Laskowski and Kato, 1980; Porten et al., 2010). On the other hand, Patient 16 presents a total of 38 *TNXB* transcripts affected by three gene variants (rs201510617, rs144415985, rs367685759) that are all predicted to be deleterious by the three scoring systems, have allele frequencies of 3.17E-02, 4.83E-02 and 5.90E-03, respectively; and in overall, are affecting two conserved domains. The first consists of a fibrinogen related domain that is present in most types of tenascins (SMART:SM00186), while the second is a fibronectin type 3 domain (SMART:SM00060) that is found in various animal protein families such as muscle proteins and extracellular-matrix molecules (Bork and Doolittle, 1992). Two of the severe patients (Patients 12 and 19) present severe-only specific compound heterozygous variants that are not predicted to be deleterious. However, one variant in the *CHGB* gene (rs742710, AF=1.07E-01), present in patient 12, has been previously reported to be potentially causative for amyotrophic lateral sclerosis early onset (Gros-Louis et al., 2009; Pampalakis et al., 2019). This gene has also been strongly suggested in literature as a possible marker for onset prediction in multiple sclerosis (Mo et al., 2013), and other related neural diseases like Parkinson’s (Nilsson et al., 2009) and Alzheimer’s disease (Chen et al., 2019). As for patient 19, the variant rs146309392 (AF=8.40E-04) in the gene *HSPG2* has been previously referred to be causal of Dyssegmental dysplasia as a compound heterozygous mutation (Arikawa-Hirasawa et al., 2001). This variant, as pointed out before, is shared with sibling patient 20. One severe individual (Patient 3), a 37 years old female, does not carry compound heterozygous variants included in this module but others at a lower resolution parameter value (**Suppl. Figure 9; Suppl. Table 6**). Interestingly, most of the genes carrying severe-specific deleterious compound heterozygous variants in this patient (*CDH3*, *FAAP100*, *FCGBP*, *GFY*, *RPTN*) are not related to processes at the NMJ level (Hull et al., 2016; Johansson et al., 2009; Kaneko-Goto et al., 2013; Ramanagoudr-Bhojappa et al., 2018; Swuec et al., 2017). Nevertheless, three of these variants occur in genes potentially involved in NMJ functionality. In particular, variants rs111709242 (AF=2.64E-03) and rs77975665 (AF=3.03E-02) affect gene *PPFIBP2*, which encodes a member of the liprin family (liprin-β) that has been described to control synapse formation and postsynaptic element development (Astigarraga et al., 2010; Bernadzki et al., 2017). Furthermore, the variant rs111709242 is predicted to be deleterious by the SIFT algorithm (see **Suppl. Table 6**). Interestingly, PPFIBP2 appears in modules at lower resolution parameter values associated with known CMS causal genes (e.g. *DOK7*, *RPSN*, *RPH3A*, *VAMP1*, *UNC13B*) (**Supplementary Figure 9)**. In addition, variant rs151154986 (AF=2.18E-02) affects the acyl-CoA thioesterase *ACOT2*, which generate CoA and free fatty acids from acyl-CoA esters in peroxisomes (Grevengoed et al., 2014). While *ACOT2* is not retained across the entire resolution range explored, community detection at the individual layer level (i.e. Louvain community detection for each network) revealed relationships with causal CMS genes at all layers (**Supplementary Figure 3**). Namely, *ACOT2* shares community membership with *ALG14, DPAGT1, GFPT1, GMPPB* and *SLC25A1*A at the protein-protein interaction layer; with *CHAT* and *SLC5A7* at the pathways layer, and with *GMPBB, SLC25A1* and *CHAT* at the metabolomic layer. A role for CoA levels in skeletal muscle for this enzyme class has been previously described (Li et al., 2015). Moreover, this patient presents high relatedness with three not-severe patients (Patients 8, 9, and 10) who in turn display a very high relatedness among them (**Suppl. Table 4**).

### Potential pharmacological implications

Finding a genetic diagnosis might help select the appropriate medication for each patient. For instance, fluoxetine and quinine are used for treating the slow-channel syndrome, an autosomal dominant type of CMS caused by mutations affecting the ligand binding or pore domains of AChR, but this treatment should be avoided in patients with fast-channel CMS (Engel et al., 2015). Within our cohort, 13 (7 mild, 2 moderate and 4 severe) out of 20 individuals from our CMS cohort are receiving a pharmacological treatment consisting of pyridostigmine, an acetylcholinesterase inhibitor used to treat muscle weakness in myasthenia gravis and CMS (Lee et al., 2018). This treatment slows down acetylcholine hydrolysis, elevating acetylcholine levels at the NMJ, which eventually extends the synaptic process duration when the AChR are mutated. Although the severity could potentially be related to how well a patient responds to the standard treatment with the AchE inhibitors, we could not find a clear correlation between severity and pyridostigmine treatment (two-tailed Fisher’s exact test p-value 0.356; **Suppl. Figure 1**). In Addition to the causal mutation in *CHRNE*, our results indicate that severity is related to AChR clustering at the Agrin-Plectin-LRP4-Laminins axis level, suggesting the potential benefit of pharmaceutical intervention enhancing the downstream process of AChR clustering. For example, beta-2 adrenergic receptor agonists like ephedrine and salbutamol have been documented as capable of enhancing AChR clustering (Clausen et al., 2018) and proved to be successful in the treatment for severe AChR deficiency syndromes (Cruz et al., 2015; Garg and Goyal, 2022). Furthermore, the addition of salbutamol in pyridostigmine treatments have been described as being able to ameliorate the possible secondary effects of pyridostigmine in the postsynaptic structure (Vanhaesebrouck et al., 2019).

### Experimental validation of USH2A involvement at the NMJ

To determine the potential relevance of one of our identified potential modifiers with no previously published relationship to the NMJ, we analyzed its function using zebrafish. For this we chose *USH2A,* a gene associated with Usher syndrome and Retinitis pigmentosa in humans (OMIM ID 608400, https://omim.org/), which was identified as a copy number loss in patient 2. While we expect the phenotypic outcome (more severe disease) of this genetic difference to manifest when expressed in conjunction with the *CHRNE* mutation causing this patients’ CMS, we hypothesized that knockdown of *USH2A* expression alone may cause detectable NMJ impairments. Therefore, we used a MO to knockdown the expression of the zebrafish orthologue; *ush2a*, and studied the effects on survival, development and NMJ function. Zebrafish *ush2a* is expressed from 1 to 5 dpf, as shown in **Suppl. Figure 10A**. Using a MO targeting the exon 3/intron 3 splice donor site we were able to decrease expression of *ush2a* with a 6 ng to 18 ng MO injection (**Suppl. Figure 10B**). Survival of control and *ush2a-* MO zebrafish was not significantly affected as compared to wildtype (WT) fish over 5 dpf (log-rank test, WT n = 574, control MO 4 ng n = 46, 6 ng n = 75, 18 ng n = 34, *ush2a-*MO 2 ng n = 72, 4 ng n = 68, 6 ng n = 360, 12 ng n = 288, 18 ng n = 139, **Suppl. Figure 10C**).

There were no obvious gross morphological differences between control MO and *ush2a-*MO fish up to 5 dpf (representative images of 2 dpf fish shown in **Suppl. Figure 10D**). As length is an indicator of developmental stage, we measured the length of 18 ng injected *ush2a-*MO fish at 2 dpf and found a significant reduction in length as compared to controls (p = 0.013, t = 2.59, df = 38, unpaired t-test, control MO n = 20, *ush2a-*MO n = 20, **Suppl. Figure 10E**). Eye area can be reduced in zebrafish models of retinitis pigmentosa, the condition that *USH2A* mutations are associated with in humans. We measured eye area in 2 dpf fish and found it to be significantly reduced in 18 ng-injected *ush2a-*MO fish as compared to controls (p = 0.0006, t = 3.73 df = 38, unpaired t-test, control MO n = 20, *ush2a-*MO n = 20, **Supplementary Figure 10F**).

Eye area remains significantly different after normalizing for body length (data not shown). CMS manifests as fatigable muscle weakness in patients and in developing zebrafish we can study the ability of fish to perform repetitive, well-characterized movements during development to determine whether impairments to the functioning of the neuromuscular system may be present. We quantified the number and duration of chorion movements in 1 dpf fish following administration of a control or 18 ng *ush2a-*MO. This revealed a significant decrease in the number of burst events performed per minute in knockdown fish as compared to controls (p = 0.003, Mann Whitney test, control MO n = 84, *ush2a-*MO n = 74, **Figure 5A**). The average duration of each burst event was not significantly affected by loss of Ush2a (p = 0.467, Mann Whitney test, control MO n = 72, *ush2a-*MO n = 49, **Figure 5B**).

**Figure 5.**
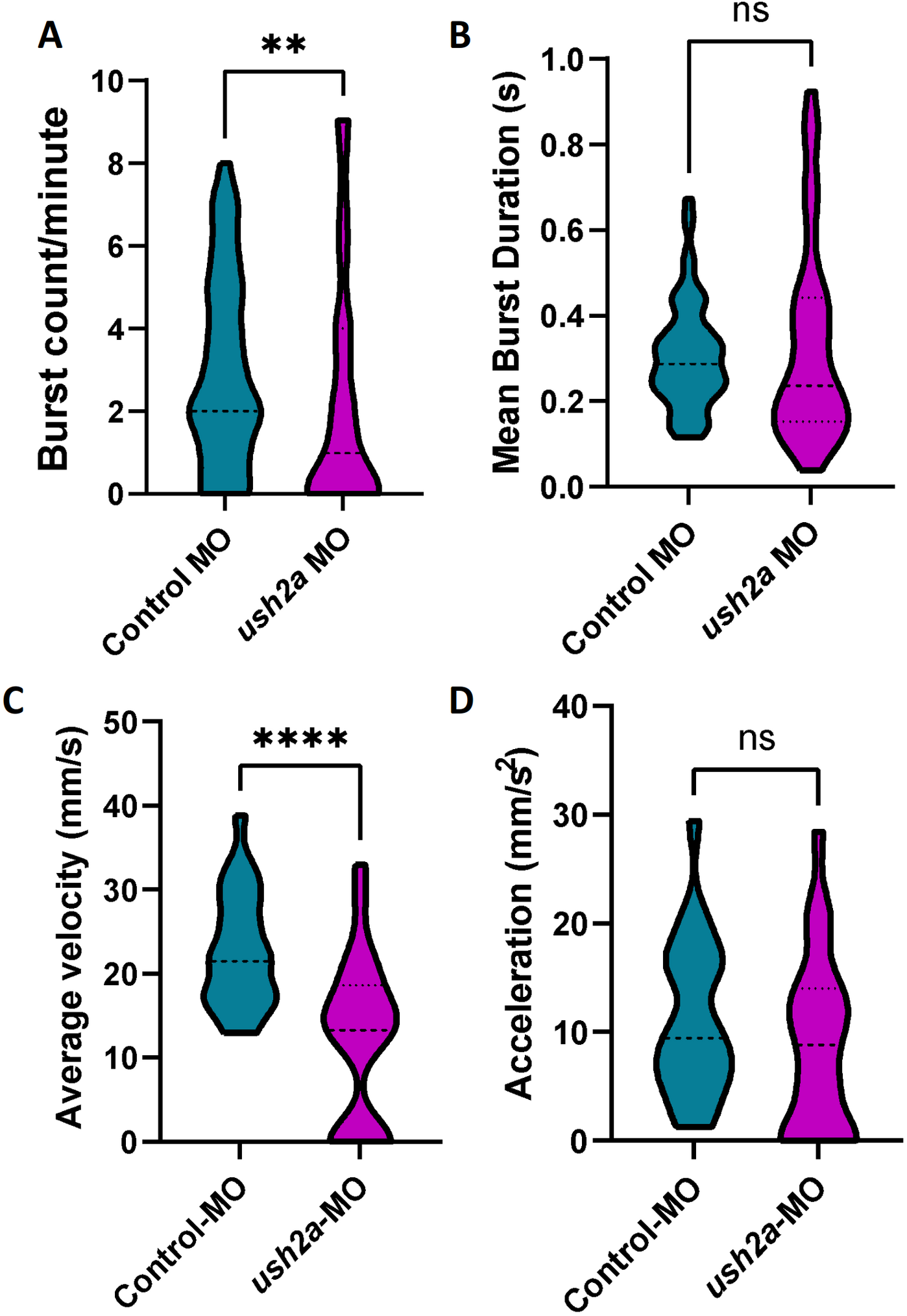
Early movement behaviors in *ush2a*-MO zebrafish. (A) Chorion rotations per minute (burst count), and (B) mean chorion rotation duration in seconds for control and *ush2a*-MO-injected zebrafish at 1 days post fertilization (dpf). (C) Average velocity and (D) initial acceleration of control and *ush2a*-MO zebrafish at 2 dpf in response to touch. Dashed line shows the median, dotted lines show the quartiles, **p < 0.01, ****p < 0.0001, ns = not significant, Mann Whitney test (A and B), unpaired t-test (C and D).

To ascertain whether impairments to movement are present in the knockdown fish while swimming free of the chorion, we also performed a touch response assay at 2 dpf. We observed a significant decrease in average velocity of the fish injected with *ush2a-*MO as compared to control MO in response to a touch stimulus (p < 0.0001, t = 4.42, df = 48, unpaired t-test; n = 25, **Figure 5C**). There was no significant difference in acceleration of *ush2a*-MO fish as compared to controls (p = 0.263, t = 1.13 df = 47, unpaired t-test; control MO n = 24, *ush2a*-MO n = 25, **Figure 5D**).

To determine whether changes in movement are reflected at the level of gross NMJ structure, analysis of NMJ morphology was performed on 2 dpf zebrafish (**Figure 6A**). A significant decrease in the number of SV2-positive clusters per 100 µm^2^ (representative of the pre-synaptic motor neurons) was identified on the fast muscle fibers of *ush2a*-MO fish as compared to controls (p = 0.0004, Mann Whitney test, control MO n = 11, *ush2a-*MO n = 15, **Figure 6B**). SV2-positive clusters overlie postsynaptic AChRs to form NMJs and these receptors can be detected with fluorophore-labelled α-bungarotoxin. Analysis of AChR clusters revealed no significant differences in number per 100 µm^2^ between the two conditions (p = 0.217, Mann Whitney test, control MO n = 11, *ush2a-*MO n = 15, **Figure 6C**). Colocalization analysis revealed no significant differences in co-occurrence of SV2 and AChR on fast muscle fibers (SV2 colocalization with AChRs: p = 0.371, t = 0.911, df = 24, nested t-test, **Figure 6D** and AChR colocalization with SV2: p = 0.372, t = 0.909, df = 24, control MO n = 11, *ush2a-*MO n = 15, nested t-test, **Figure 6E**). There was also no significant difference in colocalization of SV2 with AChRs on slow muscle, however, a significant reduction in co-occurrence of AChRs with SV2 is present on *ush2a*-MO slow muscle (SV2 colocalization with AChRs: p = 0.516, t = 0.660, df = 24, nested t-test, **Figure 6F** and AChR colocalization with SV2: p = 0.002, t = 3.41, df = 24, control MO n = 11, *ush2a-*MO n = 15, nested t-test, **Figure 6G**). Movement differences in zebrafish may also be caused by changes in muscle growth and development. Therefore, we assessed 2 dpf fish for gross phenotypic differences in muscle fiber orientation and structure using a phalloidin stain to detect actin in muscles (**Suppl. Figure 11A)**. We identified no significant differences in muscle fiber dispersion (organization) or myotome size between *ush2a*-MO and control-MO zebrafish (p = 0.922, t = 0.099, df = 24 unpaired t-test and p = 985, t = 0.019, df = 24 nested t-test, respectively. Control MO n = 11 and *ush2a-*MO n = 15, **Suppl. Figure 11B, C**).

**Figure 6.**
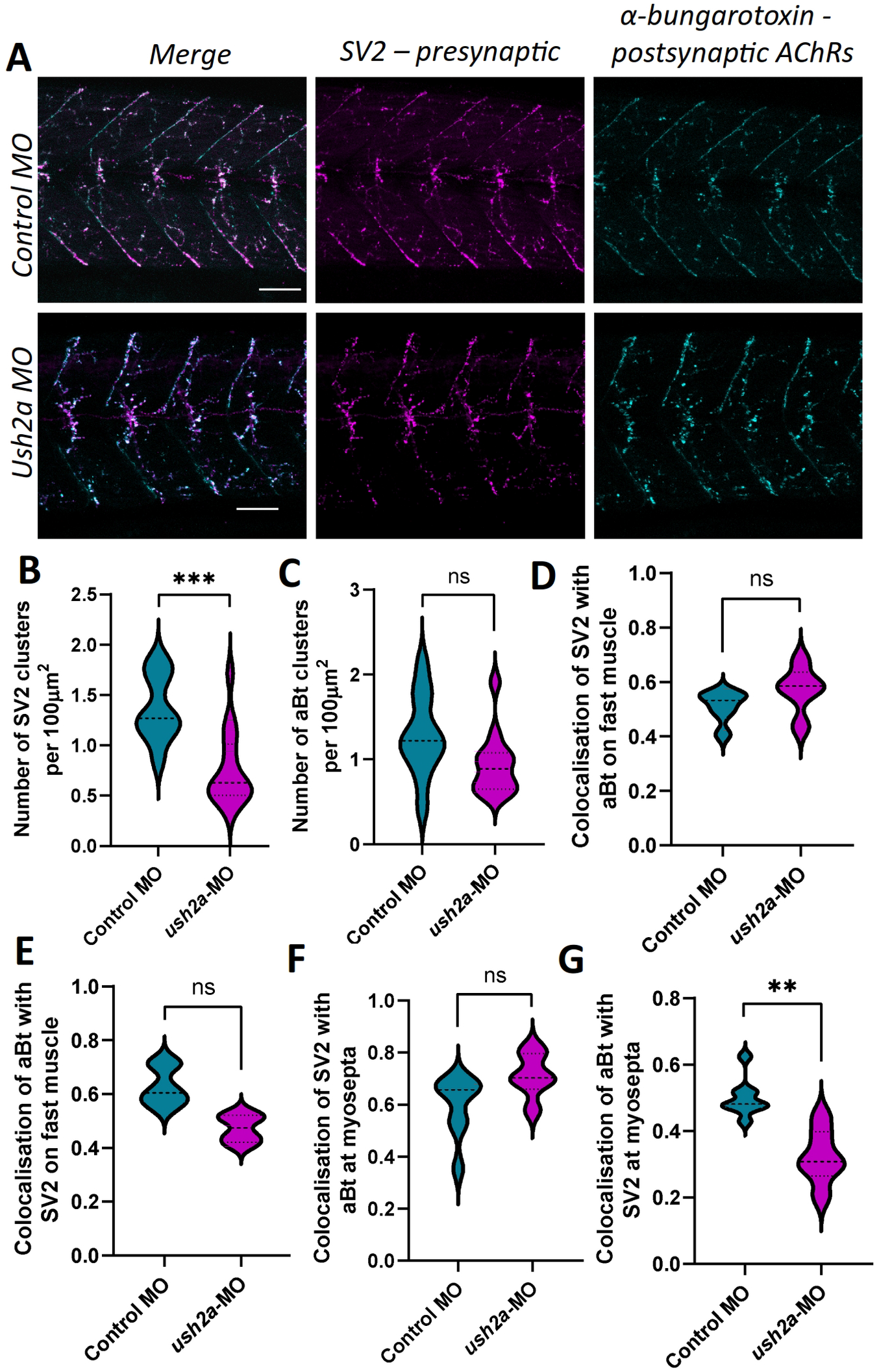
Neuromuscular junction morphology in *ush2a*-MO zebrafish. (A) Representative images of neuromuscular junctions from control and *ush2a*-MO zebrafish at 2 days post fertilization (dpf). Acetylcholine receptors (AChRs) are stained with fluorophore bound α-bungarotoxin (aBt, cyan), and motor neurons detected with an antibody against synaptic vesicle protein 2 (SV2, magenta). Scale bar = 50 µm. (B) Number of SV2-positive clusters and (C) number of aBt-positive clusters per 100 µm^2^. (D) Colocalization of SV2 with aBt and (E) colocalization of αBT with SV2 on fast muscle cells, using Mander’s correlation coefficient (0 = no colocalization, 1 = full colocalization). (F) Colocalization of SV2 with aBt and (G) colocalization of aBt with SV2 on slow muscle cells at the myosepta, using Mander’s correlation coefficient. Dashed line shows the median, dotted lines show the quartiles, **p < 0.05, ***p < 0.001, ns = not significant, nested t-test.

## Discussion

In this work, we have developed a framework for the analysis of disease severity in scenarios heavily impacted by sample size. Presenting limited numbers of cases is one of the main obstacles for the application of precision medicine methods in rare disease research, as it critically affects the level of expected statistical power, a common hallmark in the analysis of minority conditions (Whicher et al., 2018). This fact hampers exploring the molecular relationships that define the inherently heterogeneous levels of disease severity observed in rare disease populations, making it an atypically addressed biomedical problem (Boycott et al., 2013). Our approach, based on the application of multilayer networks, enable the user to account for the many interdependencies that are not properly captured by a single source of information, effectively combining the available patient genomic information with general biomedical knowledge from relevant databases representing different aspects of molecular biology. The application to a relevant clinical case, where we tested the hypothesis that the severity of CMS is determined by patient-specific alterations that impact NMJ functionality, provided evidence on how the methodology is able to recover the molecular relationships between the candidate patient-specific genomic variants, the observed causal AChR mutation and previously described CMS causal genes **(Table 1).**

Our in-depth functional analysis focused on a cohort of 20 CMS patients, from a narrow, geographically isolated and ethnically homogenous population, who share the same causative mutation in the AChR ε subunit (*CHRNE*) but present with different levels of severity. The isolation and endogamy that characterize the population from which these patients come from might have favored the accumulation of damaging variants (Fareed and Afzal, 2017; Petukhova et al., 2009), giving rise to the emergence of compound effects on relevant genes for CMS. This observation has previously been made in similar syndromes (Müller et al., 2004; Ohno et al., 2003) and in a number of other neuromuscular diseases (Wang et al., 2018; Zhong et al., 2017). Compound heterozygosity is known to happen in CMS (Hantaï et al., 2013) (Thompson et al., 2019). The initial analysis of compound heterozygous variants revealed a significant enrichment of functional categories that are specific to the severe cases, namely ECM functions. This suggests the existence of functional relationships between major actors of the NMJ that are affected by severity-associated damaging mutations. Such interactors include already known CMS causal genes (e.g. *AGRN, LRP4, PLEC*) as well as genes known to interact with them. While severity-specific compound heterozygous variants and CNVs are observed, demographic factors (e.g. sex, age), pharmacological treatment, and personalized omics data (e.g. variant calling, differential gene expression, allele specific expression, splicing isoforms) do not segregate with patient severity.

Therefore, this motivated the development of our multilayer network community analysis to investigate the relationship between known CMS causal genes and severity-associated variants (compound heterozygous variants and CNVs), integrating pathways, metabolic reactions, and protein-protein interactions. Recently, we used a multilayer network as a means to perform dimensionality reduction tasks for patient stratification in medulloblastoma, a childhood brain tumor (Núñez-Carpintero et al., 2021). Here, we started by analyzing DisGeNET data in order to verify that disease-associated genes tend to belong to the same multilayer communities. We then identified stable and significantly large gene modules within our CMS cohort’s multilayer communities and mapped the corresponding damaging mutations back to the single patients, providing a personalized mechanistic explanation of severity differences. Given the difficulties of cohort recruitment for rare diseases, this approach could be used to investigate forms of CMS and other phenotypically variable rare diseases caused by a common mutation.

Overall, our approach revealed major relationships at the protein-protein and pathway layers. The personalized analysis of these mutations further suggests that CMS severity can be ascribed to the damage of specific molecular functions of the NMJ which involve genes belonging to distinct classes and localizations, namely ECM components (proteoglycans, tenascins, chromogranins) and postsynaptic modulators of AChR clustering (*LRP4, PLEC*) (**Table 2**). Alterations of other genes related to ECM components, such as laminins and collagen, are observed but are not specific to the severity levels.

Although at first the use of metabolomic knowledge in the multilayer network did not seem to provide highly relevant information for the cohort, it provided relevant insights for the personalized analysis of patient 3, whose mutations presented functional relationships in all layers with other CMS causal genes outside of the presented severe-specific module (**Supplementary Figure 3**).

Finding a personalized genetic diagnosis for phenotypic severity might help select the appropriate medication for each patient. For instance, fluoxetine and quinidine are used for treating the slow-channel syndrome, an autosomal dominant type of CMS caused by mutations affecting the ligand binding or pore domains of AChR, but this treatment should be avoided in patients with fast-channel CMS (Engel et al., 2015). Within our cohort, 13 out of 20 individuals from our CMS cohort are receiving a pharmacological treatment consisting of pyridostigmine, an acetylcholinesterase inhibitor used to treat muscle weakness in myasthenia gravis and CMS (Lee et al., 2018). Although the severity could potentially be related to how well a patient responds to the standard treatment with the AchE inhibitors, we could not find a clear correlation between severity and pyridostigmine treatment (two-tailed Fisher’s exact test p-value 0.356; Suppl. Figure 1). Our results indicate that severity is related to AChR clustering at the Agrin-Plectin-LRP4-Laminins axis level, suggesting the potential benefit of pharmaceutical intervention enhancing the downstream process of AChR clustering. Strikingly, beta-2 adrenergic receptor agonists like ephedrine and salbutamol have been documented as capable of enhancing AChR clustering (Clausen et al., 2018) and proved to be successful in the treatment for severe AChR deficiency syndromes (Rodríguez Cruz et al., 2015; Garg and Goyal, 2022; Sadeh et al., 2011; Vanhaesebrouck et al., 2019), but a strong molecular explanation for the observed favorable effects was still missing. This study reinforces explainability for the described successful usage of such treatments by relating CMS phenotypic severity with the normal development of AChR clusters at the motor neuron membrane.

Several of the genes identified in this analysis do not have previous associations with the NMJ, such as the Usher syndrome and Retinitis pigmentosa associated gene; USH2a, identified as a copy number loss in patient 2. To provide proof of principal for this gene acting as a potential modifier of CMS severity, we investigated whether knockdown of ush2a, the zebrafish orthologue, could result in NMJ defects. Both CRISPR and TALEN-mediated knockout of ush2a in zebrafish have previously revealed phenotypes consistent with Usher syndrome and Retinitis pigmentosa such as hearing loss and progressive visual impairments (Han et al., 2018). However, neither study assessed impacts on muscle structure or movement of the fish. Zebrafish perform well-characterized movements throughout development, starting with spontaneous chorion rotations from approximately 17 hours post fertilization (hpf, the time at which primary motor axons start extending into the muscle) to 30 hp (Saint-Amant and Drapeau, 1998). We treated 1-cell-stage embryos with a high dose of MO to reduce expression of ush2a (or equivalent dose of a control MO) and found a decrease in the number of chorion rotations performed at 24 hpf. These movements are mediated at the level of the spinal cord and are independent of supraspinal inputs (Downes and Granato, 2006), thus implying an early defect in NMJ or muscle development, or in signal transduction in the spinal cord/peripheral nervous system. By 2 dpf zebrafish can respond to touch and do so by rapidly swimming at least 1 body-length away from the stimulus (Saint-Amant and Drapeau, 1998). In ush2a-MO fish the average swimming velocity was significantly slower than in controls, whereas the initial acceleration (proportional to the force of muscle contraction) was unaffected (Sztal et al., 2016). This implies that the initial fast muscle response is not significantly affected at this time-point, but that loss of Ush2a at the NMJs of slow muscle may be impacting swimming. Defects in movement are reported in many other zebrafish models of CMS, such as those lacking Dok7 (Müller et al., 2010), Gfpt1 (Senderek et al., 2011) and Syt2 (Wen et al., 2010). Our motility findings are supported by the identification of a reduction in colocalization of AChRs with SV2-positive clusters on slow muscle fibers in 2 dpf fish, thus showing an increase in the number of AChRs that have not been contacted by a motor axon. We also identified an overall reduction in the number of SV2-positive clusters, which may be indicative of a defect or delay in development of the motor nervous system. Previous studies have commented on USH2A presence on the basement membranes of perineurium nerve fibers (Pearsall et al., 2002; Schwaller et al., 2021), however, further studies in a mammalian model and/or using zebrafish mutants rather than transient knockdown will be required to determine the presence of USH2a at the NMJ, and whether loss of USH2a alone can impact NMJ signaling or whether co-occurrence with CHRNE CMS is required. Additional functional work is also required to ascertain the importance of other potential modifiers identified in this study. Particularly, a prospective analysis on the potential NMJ involvement of the unique variants detected for the non-severe group could be of special interest for the study of CMS, potentially discerning their functional relationship to causal CMS genes.

Our work represents a thorough study of a narrow population showing a differential accumulation of damaging mutations in patients with CMS who have varying phenotypic severities, building on the initial impact of *CHRNE* mutations on the NMJ. It is important to remark that CMS is of particular interest among rare diseases, since drugs that influence neuromuscular transmission can produce clear improvements in the affected patients (Engel, 2007). In this sense, identifying meaningful molecular relationships between gene variants allow us to gain insight into the disease mechanisms through a multiplex biomedical framework, paving the way for a whole new set of computational approximations for rare disease research.

## Acknowledgments

The authors acknowledge the donors and families, Daniel Rico (Newcastle University) for his contribution in early stages of the project, Anaïs Baudot (Aix Marseille Université and Barcelona Supercomputing Center) for her careful revision of the manuscript, Miguel Vázquez (Barcelona Supercomputing Center) for advising about Rbbt analysis, Jon Sánchez-Valle (Barcelona Supercomputing Center) and Núria Olvera (Barcelona Supercomputing Center and IDIBAPS) for the insightful discussions.

## Author contributions

T.C., I.T. and V.G. collected and processed the biopsies; H.L. and R.T. coordinated data sharing; A.T., P.A.C.T., S.B. and S.C. coordinated and performed the omics data analysis with Y.A., S.L., M.R. and M.B.; E.O. and S.S. performed the experimental validations; D.C. and A.V. coordinated the multilayer network analysis performed by I.N.C. All authors contributed to the writing and revising of the manuscript.

## Ethics approval

This study was approved by the Ethics committee of Sofia Medical University (protocol 4/15-April-2013). Moreover, we have complied with all relevant ethical regulations regarding animal research.

## Methods

### WGS and RNA-seq

Whole genome sequencing (WGS) data have been obtained from blood using the Illumina TruSeq PCR-free library preparation kit. Sample sequencing was performed with the HiSeqX sequencing platform (HiseqX v1 or v2 SBS kit, 2×150 cycles), with an average mean depth coverage ≥ 30X. Samples have been analyzed using the RD-Connect specific pipeline: BWA-mem for alignment; Picard for duplicate marking and GATK 3.6.0 for variant calling. RNA sequencing (RNA-seq) data have been obtained from fibroblasts, using Illumina TruSeq RNA Library Preparation Kit v2, sequencing with an average of 60M reads per sample (paired-end 2X125 cycles). Data has been processed with the following pipeline (Laurie et al., 2016): STAR 2.35a for alignment, RSEM 1.3.0 for quantification, and GATK 3.6.0 for variant calling. All analyses have been performed using the human genome GRCh37d5 as reference.

### Copy number variants

Copy Number Variants (CNVs) have been extracted using ClinCNV (https://github.com/imgag/ClinCNV) by employing a set of Eastern European samples as a background control group. Out of the 569 autosomal CNVs we selected as potential candidates the CNVs of the following types that overlapped with protein-coding genes: 1) whole gene gains or losses, and 2) partial losses (deletions overlapping with exons but not with the whole gene). The list of potential candidates included 55 CNVs that created a total of 82 whole gene gains or losses and 28 partial losses.

### Compound heterozygous variants

Compound heterozygous variants have been obtained by phasing the WGS variant calls with the RNA-seq aligned BAM files using phASER (Castel et al., 2016). At first, variants are imputed using Sanger Imputation Service with EAGLE2 pre-phasing step (Durbin, 2014). PhASER is then applied to extend phased regions to gene-wide haplotypes. By accurately reflecting the muscle transcriptome, fibroblasts have been previously proved to be excellent and minimally invasive diagnostic tools for rare neuromuscular diseases (Gonorazky et al., 2019).

We then annotated variants with eDiVA tool (www.ediva.crg.es) (Bosio et al., 2019), and removed all mutations with Genome Aggregation Database (gnomAD) (Lek et al., 2016) that show allele frequency > 3% globally, all variants outside exonic and splicing regions using Ensembl annotation, all synonymous mutations, and all variants with read depth (coverage) smaller than 8. Afterwards we selected all genes with at least two hits on different alleles as genes affected by damaging compound heterozygous variants. Each sample has been processed individually throughout the whole process.

### Monolayer community detection

We performed a network community detection analysis using the Louvain clustering algorithm (Blondel et al., 2008) implemented in R package igraph (https://igraph.org/) with default parameters. We carried out the analysis using three (monolayer) networks, obtained from Reactome database (Fabregat et al., 2018), from the Recon3D Virtual Metabolic Human database (Brunk et al., 2018) (both downloaded in May 2018), and from the Integrated Interaction Database (IID) (Kotlyar et al., 2019) (downloaded in October 2018). Additional information on network connectivity metrics (e.g. node centrality distributions and specific centrality information for severe-specific module genes) is conveniently provided as a Jupyter Notebook, accesible at the following link: https://github.com/ikernunezca/CMS/blob/master/Scripts/Multilayer_Network_Information_and_Connectivity_Patterns.ipynb.

All gene identifiers of each network were converted to NCBI Entrez gene identifiers using R packages AnnotationDbi v1.44.0 and org.Hs.eg.db v3.7.0 (http://bioconductor.org/). After detecting the community structure from each layer independently, we retrieved the community membership of the genes of interest, henceforth called “CMS linked genes”, i.e. known CMS causal genes, and severe and not-severe compound heterozygous variants and CNVs. We then defined a community similarity measure as Jaccard Index, i.e. the number of shared genes of interest between the communities divided by the sum of the total number of genes of each community.

### Multilayer community detection

We constructed a multilayer gene network composed of the three monolayer networks described in the previous section (Reactome, Virtual Metabolic Human and Integrated Interaction Database). Each of these three networks represents one layer of the multilayer network and, in general, three facets of fundamental molecular processes in the cell (**Suppl. Figure 11**). The multilayer community detection analysis was performed using MolTi software (Didier et al., 2015), which adapts the Louvain clustering algorithm with modularity maximization to multilayer networks. The algorithm is parametrized by the resolution (γ): the higher the value of γ, the smaller the size of the detected multilayer communities. By varying the resolution parameter γ it is possible to uncover the modular structure of network communities (Fortunato and Barthélemy, 2007).

By exploring a wide range of resolution parameter values, we identified γ=4 (727 communities, each one composed of 26.46 genes on average) as an extreme value before both size and number of the detected multilayer communities stabilize (**Suppl. Figure 12**). The most dramatic changes in number and composition of detected communities are observed in the resolution parameter interval γ∈(0,4].

We, therefore, used this parameter interval to test the hypothesis that disease-related genes consistently appear in the same multilayer communities, as well as to identify modules containing CMS linked genes within them. In this analysis, we define a module as a group of CMS linked genes that are systematically found to be part of the same multilayer community while increasing the resolution parameter (see Supplementary Information "Multilayer community detection analysis").

### Additional analyses and code availability

We retrieved known CMS causal genes from the GeneTable of Neuromuscular Disorders (http://www.musclegenetable.fr, version November 2018) (Bonne et al., 2017). Segregation analysis of WGS data has been performed using Rbbt (Vázquez et al., 2010). DisGeNET database (Piñero et al., 2017) was downloaded in November 2018. The association between CMS severity, demographic factors and clinical tests was assessed with a two-tailed Fisher’s test using R statistical environment (www.R-project.org). Networks were rendered with Cytoscape (Saito et al., 2012). We used VCFtools (Danecek et al., 2011) to compute familial relatedness Ω among patients, scaled to -log_2_(2Ω). We used Enrichr (Chen et al., 2013) for the functional enrichment analysis of the gene lists under study. We used Ensembl Variant Effect Predictor (VEP)(McLaren et al., 2016) to assess the impact of the compound heterozygous variants in the genes of the severe-specific largest module. Expression levels in tissues of interest (GTEx and Illumina Body Map) were retrieved from EBI Expression Atlas (www.ebi.ac.uk/) by filtering with the following keywords: ‘nerve’, ‘muscle cell’, ‘fibroblast’ and ‘nervous system’ (0.5 TPM default cutoff). We used Expression Atlas expression level categories: low (0.5 to 10 TPM), medium (11 to 1000 TPM), and high (more than 1000 TPM) (Petryszak et al. 2016). Synaptic localization was retrieved from the UniProt database (https://www.uniprot.org/).

### Zebrafish morpholino injections

Zebrafish have one orthologue of human *USH2a: ush2a*, as identified using the UCSC database (http://genome.ucsc.edu/, GRCz11/danRer11 assembly). We confirmed that *ush2a* is expressed throughout the first 5 days post fertilization (dpf). Gene Tools LLC (USA) then designed and synthesized an antisense morpholino oligonucleotide (MO) targeting the splice donor site of exon 3/intron 3 of *ush2a* (5’-3’ GAGAAATGCTGCTCACCTGTAGAGC, ENSDART00000086201.5). We also obtained a control MO that targets a human beta-globin mutation (5’-3’ CCTCTTACCTCAGTTACAATTTATA). MOs were diluted to 2 ng/nl in Danieau buffer (58 mM NaCl, 5 mM HEPES, 0.7 mM KCl, 0.6 mM Ca(NO_3_)_2_, 0.4 mM MgSO_4_; pH 7.6) and supplemented with 1% phenol red, before being injected into the yolk-sac of 1-cell stage embryos. A range of doses between 6 and 18 ng per 1-cell stage embryo were trialed for success in reducing *ush2a* expression and producing a measurable phenotypic change. A dose of 18 ng per 1-cell stage embryo was selected for behavioral and morphological analysis, as survival was not significantly affected for any dose tested. Embryos were maintained at 28.5°C in blue water (system water with 0.1 µg/ml Methylene Blue) for up to 5 dpf and survival recorded daily. At 2 dpf zebrafish were imaged using a Leica EZ4 W stereomicroscope and eye size and length measured using Fiji (ImageJ).

### Chorion movement analysis in zebrafish

At 1 dpf (24 hours post fertilization), zebrafish were recorded in their chorions for 1 minute at 30 frames per second using a Leica EZ4 W stereomicroscope. Videos were analyzed using DanioScope software (Noldus Information Technology Inc., Leesburg, VA) to automatically assess duration of bursts and burst count/minute (bursts are full rotations performed by the zebrafish within the chorion).

### Touch response analysis

At 2 dpf, a touch response assay was performed as previously described (O’Connor et al., 2018). Only fish with a normal phenotype were used for movement analysis. Briefly, fish that had not hatched from the chorion were enzymatically dechorionated with pronase (1 mg/ml, Sigma) for 10 min in blue water, followed by 3x washes in blue water. An individual fish was placed in a petri dish containing blue water and a Sony RX0 II (DSC-RX0M2) camera was placed 20 cm above the petri dish. A ruler with 1 mm markings was used as a scale for recordings. A gel loading pipette tip was used to touch the zebrafish on the back of the head and the response recorded. Videos were imported into Fiji ImageJ (Schindelin et al., 2012) as FFmpeg movies and movements analyzed using the Trackmate plugin (Tinevez et al., 2017). Values for average speed were exported and used to derive initial acceleration.

### RNA isolation, cDNA synthesis and RT-PCR in zebrafish

RNA was isolated from pools of around 20 2 dpf zebrafish (control MO and *ush2a* MO-injected) following removal of chorions with pronase (Streptomyces griseus, Roche,1 mg/ml in blue water). Zebrafish were washed 3 times with blue water, euthanized with a 1:1 ratio of fresh system water:4 mg/ml tricaine methanesulfonate (Sigma). Fish were homogenized in RLT buffer (RNeasy mini kit, Qiagen) using 5 mm stainless steel beads with a TissueLyser II (Qiagen) at 25 Hz for 2 mins. RNA was then isolated following the RNeasy kit manufacturer’s instructions, including on-column DNase digestion. RNA was measured using a Nanodrop ND-1000 and 1 µg used for cDNA synthesis according to manufacturer’s instructions (5X All-In-One RT MasterMix, abm). Reverse-transcriptase PCR (RT-PCR) was performed to check for *ush2a* gene expression and knockdown success in MO-treated embryos, using MyTaq™ DNA Polymerase (Meridian Bioscience) and primers as follows: *eef1a1l1* forward 5’-CTGGAGGCCAGCTCAAACATGG-3’, reverse 5’-CTTGCTGTCTCCAGCCACATTAC-3’ and *ush2a* forward 5’-CTGGGCACACTTGGCTCTAC -3’, reverse 5’-TTCTTCAATCTCCCTGTTGGTT-3’.

### Immunofluorescent staining, imaging and analysis of zebrafish neuromuscular junctions and muscle fibers

Whole mount staining of 2 dpf zebrafish NMJs was performed as previously described (O’Connor et al. 2019). Briefly, a mouse anti-synaptic vesicle protein 2 (SV2) antibody was used to visualize motor neurons (1:200, AB2315387, Developmental Studies Hybridoma Bank) and Alexa Fluor 488-α-bungarotoxin conjugate (1:1000, B13422, Invitrogen) was used for visualizing acetylcholine receptors (AChRs). Phalloidin-iFluor 594 was used to visualize filamentous actin within muscle fibers 1:1000, ab176757). Z-stack images encompassing the depth of the midsection of the zebrafish tail were obtained using a 20× air objective on an LSM800 confocal microscope. Analysis of NMJ structure was performed as previously described (O’Connor et al., 2019), using Fiji (ImageJ, Madison, WI, USA). The number of SV2-positive and α-bungarotoxin-positive clusters per 100 µm^2^ were measured. Co-localization analysis between SV2 and α-bungarotoxin was performed on maximum intensity projections using the ‘JACoP’ Fiji plugin (Bolte and Cordelières, 2006). Briefly, each fluorophore was subject to manual thresholding to remove background, and the Mander’s correlation coefficient calculated to give a value between 0 and 1, reflecting the degree of co-occurrence of signals between both SV2 and α-bungarotoxin, and α-bungarotoxin with SV2. For phalloidin-stained fish, average myotome size was measured, and degree of fiber dispersion quantified using the directionality plugin. Data was collected from at least 4 myotomes per fish.

### Statistics for zebrafish experiments

Statistical analysis was performed using GraphPad Prism software (v9.3.0). Outliers were removed from data using the ROUT method (Q = 1 %). Cleaned data was tested for normal distribution then depending on outcome either a nonparametric Mann-Whitney test or parametric unpaired t-test were applied for behavioral studies and degree of dispersion. For NMJ morphology experiments in which 4+ myotomes (technical replicates) per fish (biological replicates) were analyzed, data was assessed for significance using a nested t-test to avoid pseudoreplication. Statistical significance was taken as p < 0.05, degrees of freedom (df) and t-value are given for all parametric tests, and n numbers listed in the results section. Survival analysis was performed using the log-rank test comparing WT to each other condition, and threshold for significance was corrected for multiple comparisons using the Bonferroni method (p < 0.006). Zebrafish studies were blinded before image/video acquisition and unblinded following analysis.

### Data availability

The datasets generated and analyzed in this study are not publicly available due to sensible content (genomics information in a rare disease). Reasonable requests for further information will be carefully evaluated by the corresponding author and co-authors.

### Code availability

All code and the Cytoscape session rendering Figures 3 and 4, as well as Supplementary Figures 3, 6 and 9 are available for reproducibility purposes at: https://github.com/ikernunezca/CMS. The analysis of multilayer community communties can also be performed using CmmD (Núñez-Carpintero et al., 2021) (https://github.com/ikernunezca/CmmD) with parameters: resolution_start: 0, resolution_end: 4, interval: 0.5 and the CMS linked genes as nodelist.

