## Supplementary Figures and Supplementary Information for "Rare disease research workflow using multilayer networks elucidates the molecular determinants of severity in Congenital Myasthenic Syndromes"

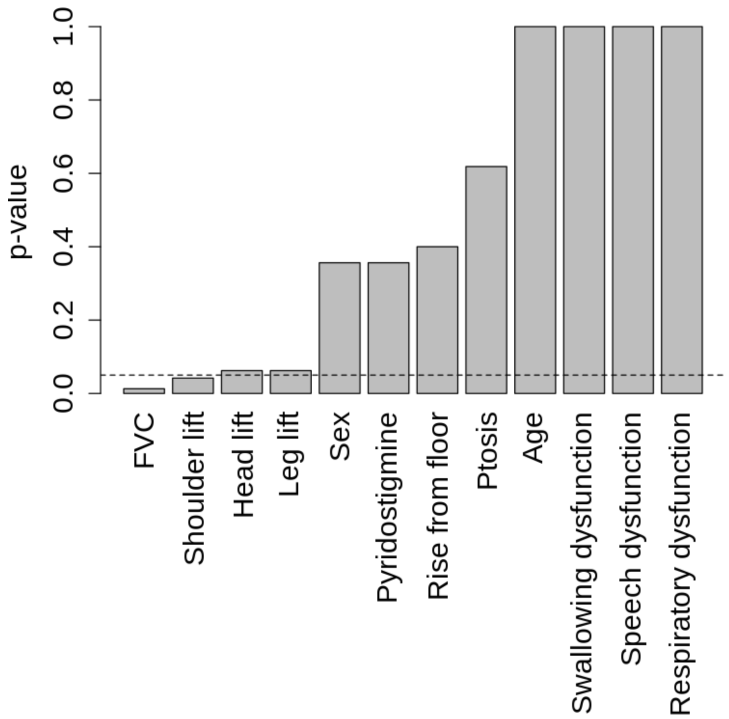


**Suppl. Figure 1.** Association between CMS severity (severe and not-severe phenotypes) and demographic factors (age, sex), pharmacological treatment (pyridostigmine), and clinical tests (speech, respiratory, swallowing functionality, ability to list shoulder, head, leg, eyelids (ptosis), and to rise from the floor, and Forced Vital Capacity (FVC)) (Suppl. Table 1). Classes were defined based on Suppl. Table 1. Age was discretized into two classes ('young' and 'old') based on the average age of all the patients (40 years). Barplot reports the p-values of a two-tailed Fisher's exact test (Methods). The dotted line indicates a p-value of 0.05.

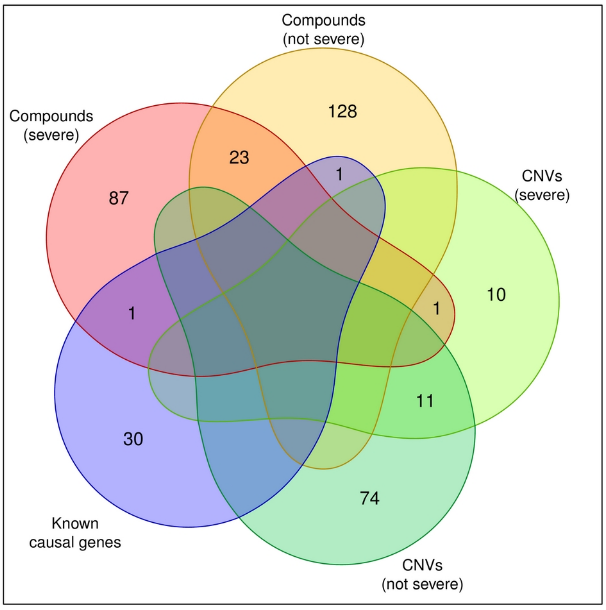


**Suppl. Figure 2.** Venn diagram of the genes associated with CNVs and compound heterozygous variants in not-severe and severe phenotypes as well as known CMS causal genes.

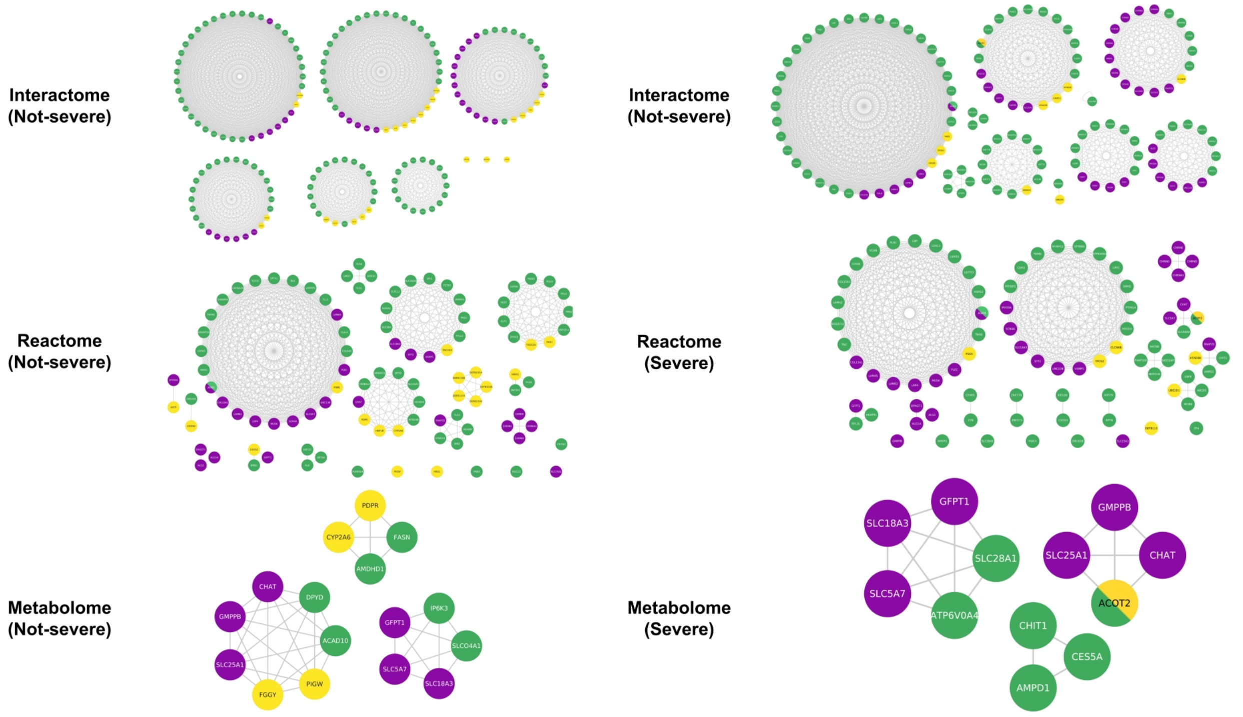


**Suppl. Figure 3.** Communities of CMS linked genes in the monolayer networks. Nodes are connected if they share membership to the same community from the clustering obtained using the Louvain algorithm. In green compound heterozygous variants; in yellow, CNVs; in purple, known CMS causal genes. Being a causal gene bearing compound heterozygous variants, AGRN is depicted in both purple and green. Being a gene presenting both compound heterozygous mutations and copy number variations, *ACOT2* is depicted in both green and yellow.

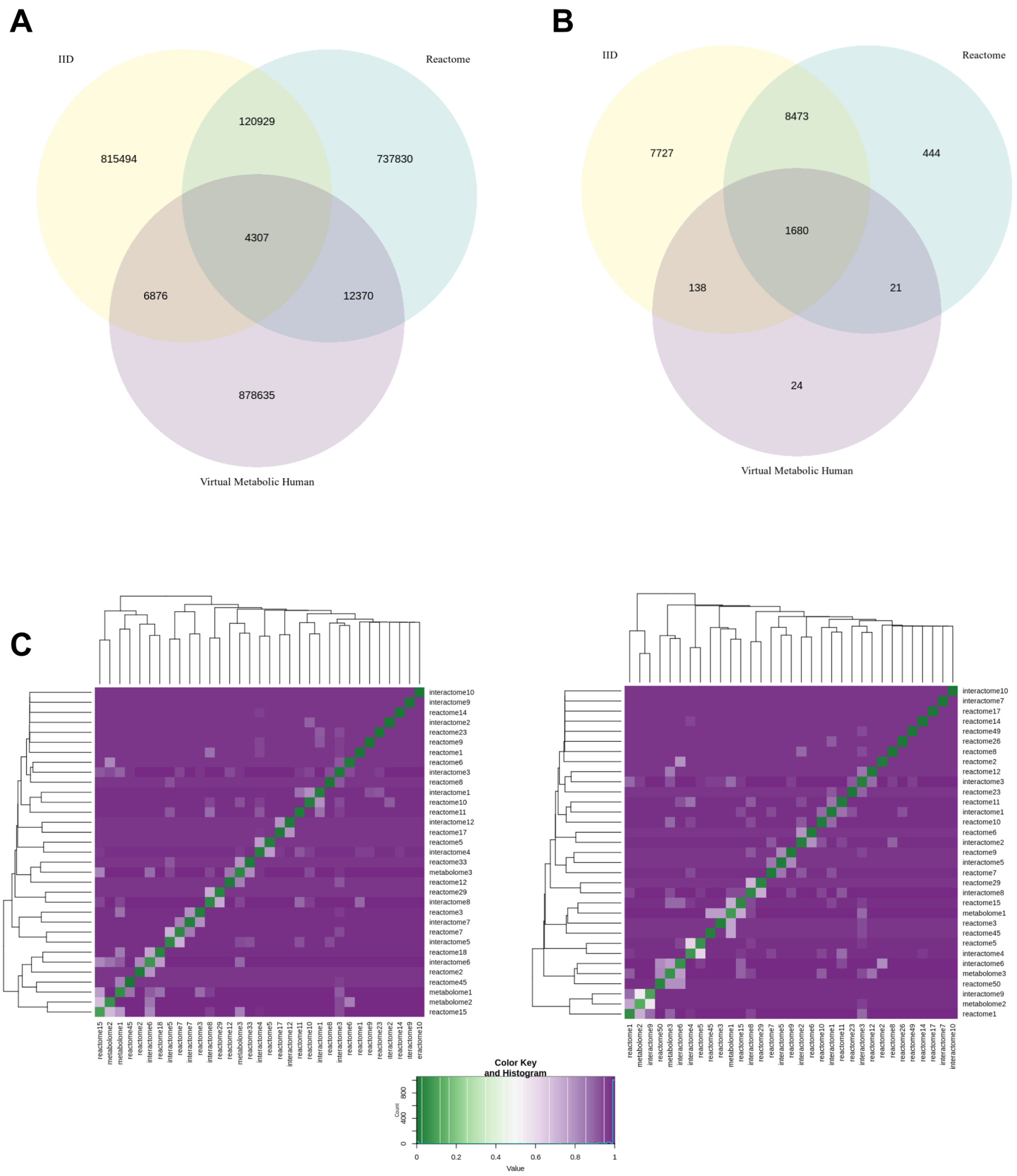


**Suppl. Figure 4.** (**A**) Edge overlap among the layers of the multilayer network. Each layer is identified by the name of the database from which the information was retrieved. (**B**) Node overlap among the layers of the multilayer network. (**C**) Heatmap of the Jaccard index dissimilarity among communities of CMS linked genes in the monolayer networks. Left, not-severe group, right, severe group.

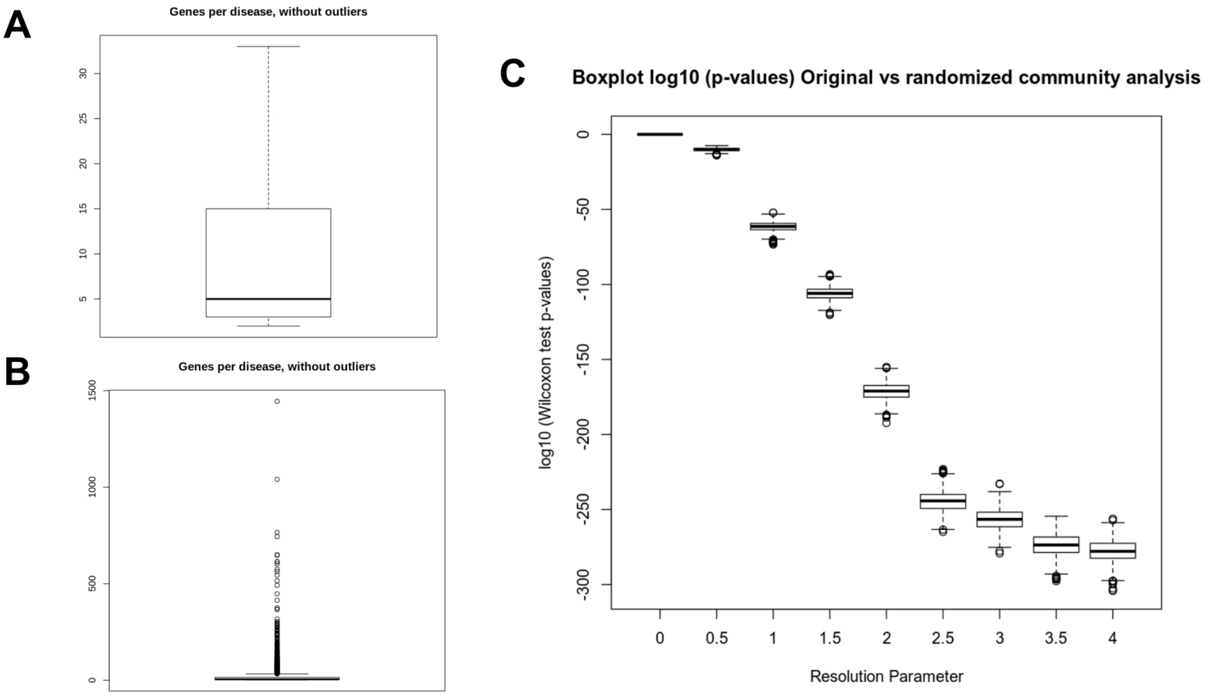


**Suppl. Figure 5**. Distribution of the number of genes per disease in the DisGeNET database, not showing (**A**) and showing (**B**) outliers (Methods). Distribution of p-values (two-sided Wilcoxon test; logarithmic scale) associated with DisGeNET multilayer communities along the range of the MolTi resolution parameter under evaluation (**C**; Methods).

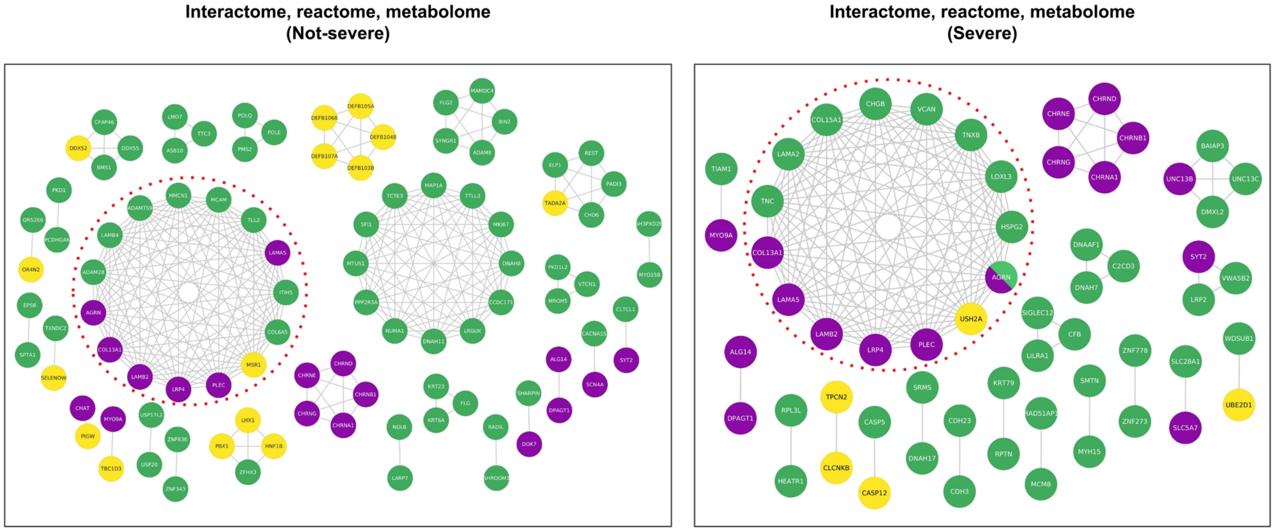


**Suppl. Figure 6.** Multilayer modules containing CMS linked genes of the severe and not-severe groups. Nodes are connected if the genes share membership to the same multilayer community across the range of MolTi resolution parameter (Methods). Modules with a size not expected to be found by chance are highlighted with a dotted circle (p-value < 0.05). In green compound heterozygous variants; in yellow, CNVs; in purple, known CMS causal genes.

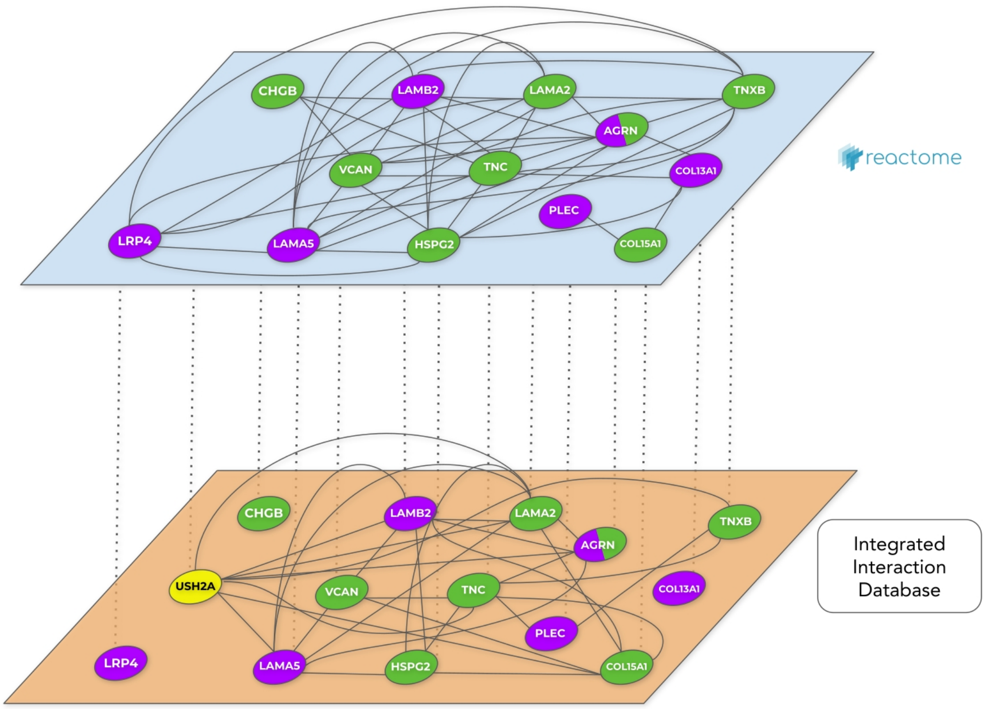


**Suppl. Figure 7.** Nature of the existing incident interactions between the genes identified in the severe-specific module. In green compound heterozygous variants; in yellow, CNVs; in purple, known CMS causal genes. As *LOXL3* do not present incident interactions in any of the two layers but with other module genes that are not represented for not being a CMS linked gene, *LOXL3* is not depicted. Provided that *USH2A* do not exist on the pathways layer, it is only depicted on the protein-protein interaction layer.

Tissue Expression for AGRN (TPM)

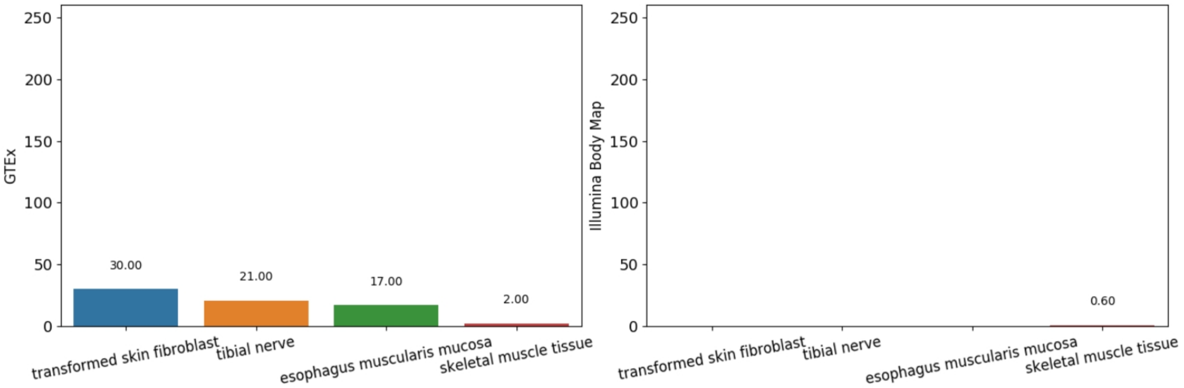


Tissue Expression for CHGB (TPM)

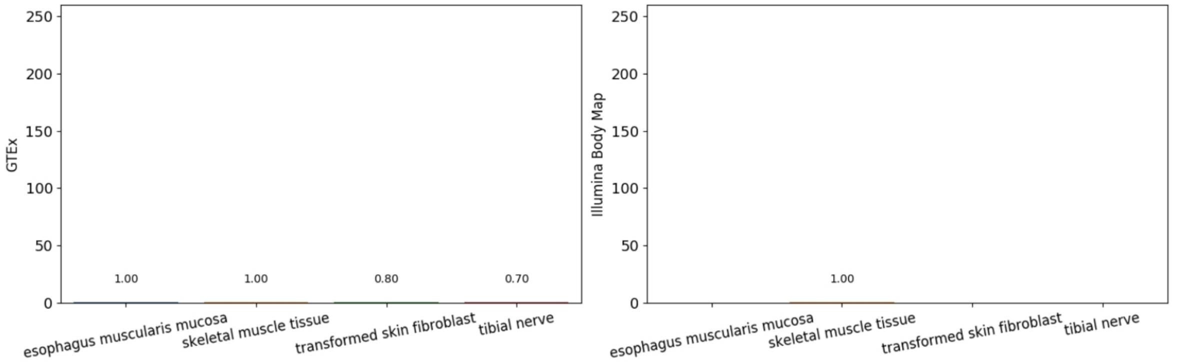


Tissue Expression for COL13A1 (TPM)

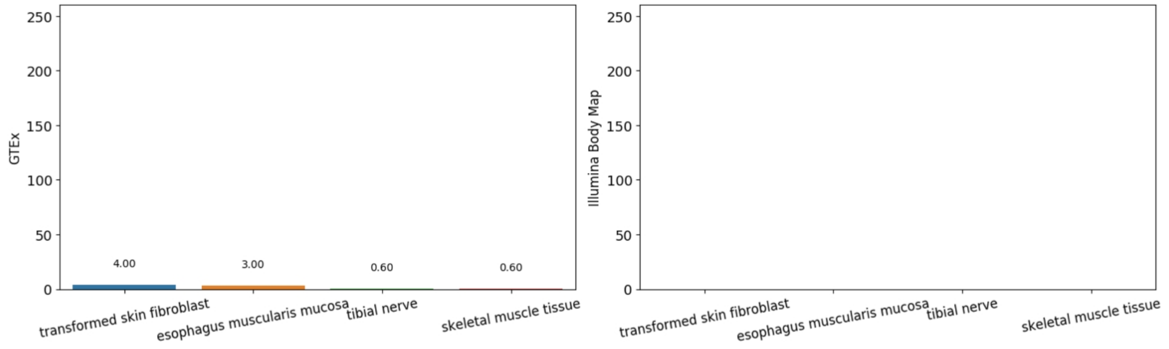


Tissue Expression for COL15A1 (TPM)

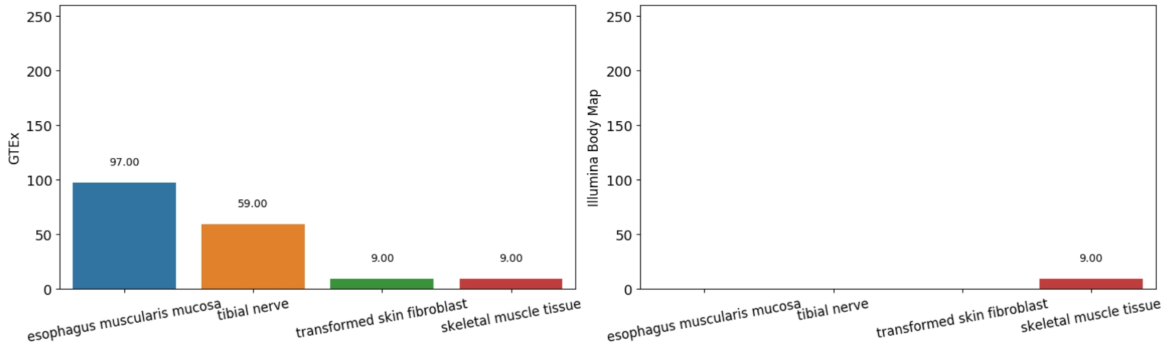

Tissue Expression for HSPG2 (TPM)

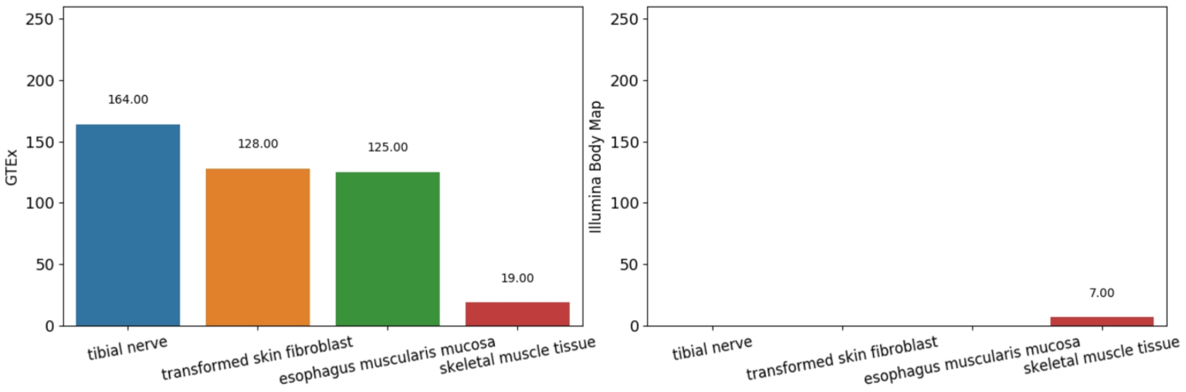


Tissue Expression for LAMA2 (TPM)

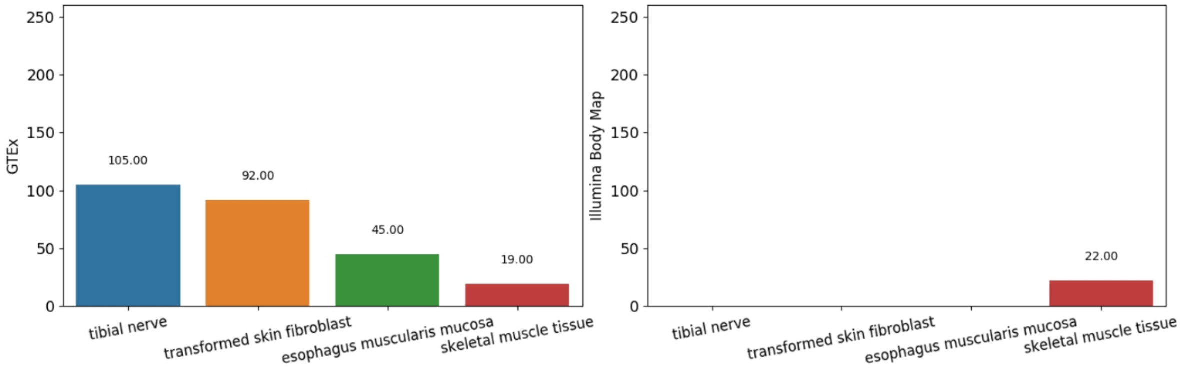


Tissue Expression for LAMA5 (TPM)

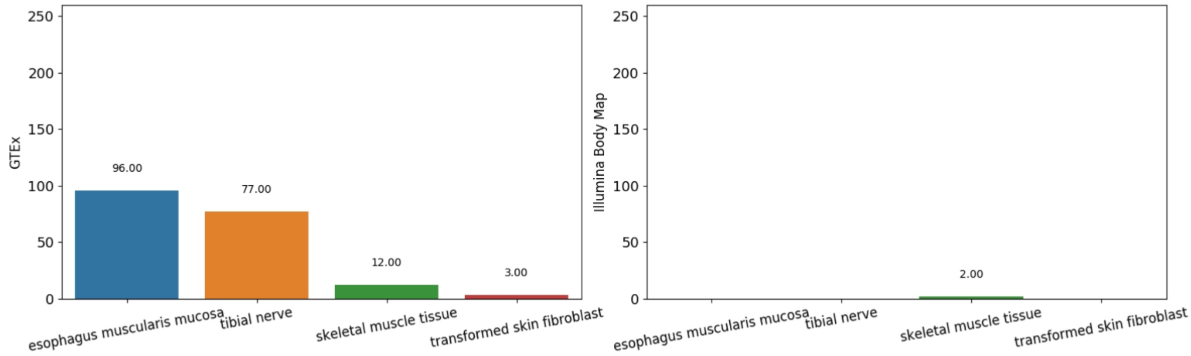


Tissue Expression for LAMB2 (TPM)

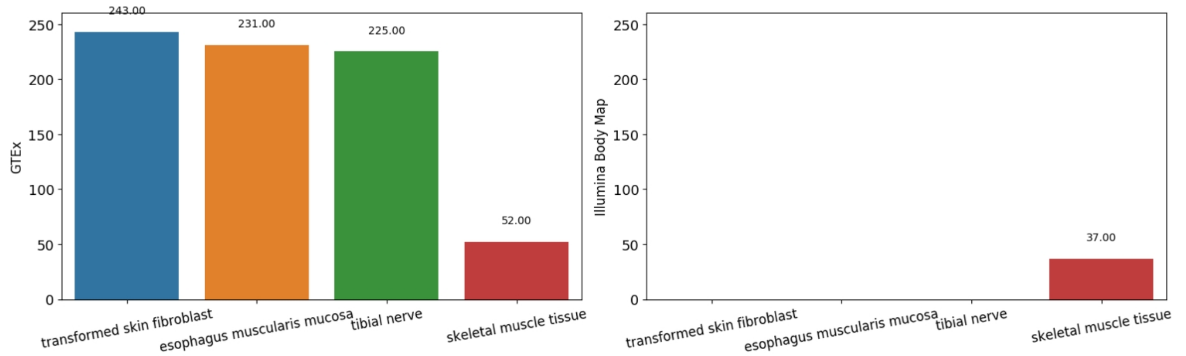

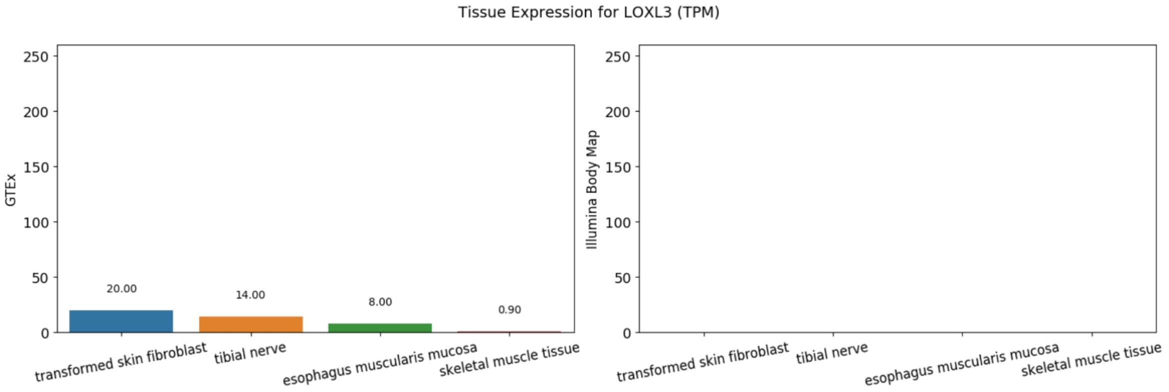
Tissue Expression for LOXL3 (TPM)
GTEx
Illumina Body Map
0
50
100
150
200
250
20.00
14.00
8.00
0.90
transformed skin fibroblast
tibial nerve
esophagus muscularis mucosa
skeletal muscle tissue

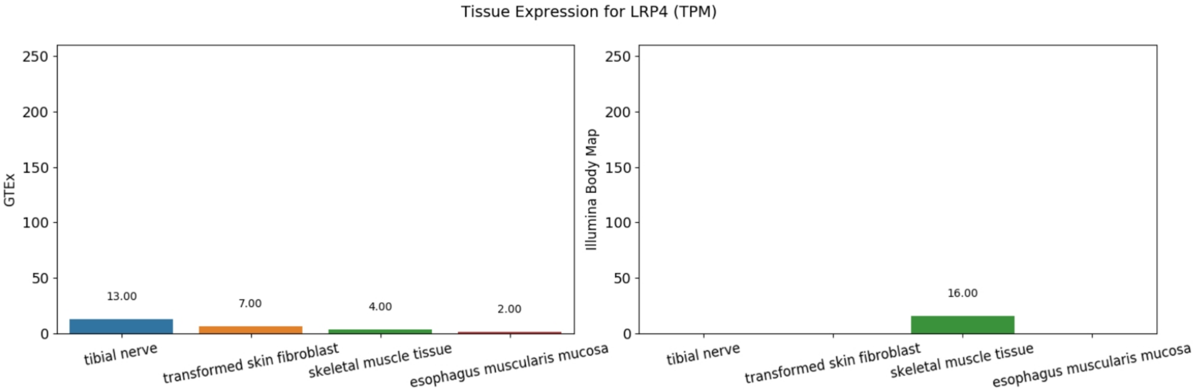
Tissue Expression for LRP4 (TPM)
GTEx
Illumina Body Map
0
50
100
150
200
250
13.00
7.00
4.00
2.00
16.00
tibial nerve
transformed skin fibroblast
skeletal muscle tissue
esophagus muscularis mucosa

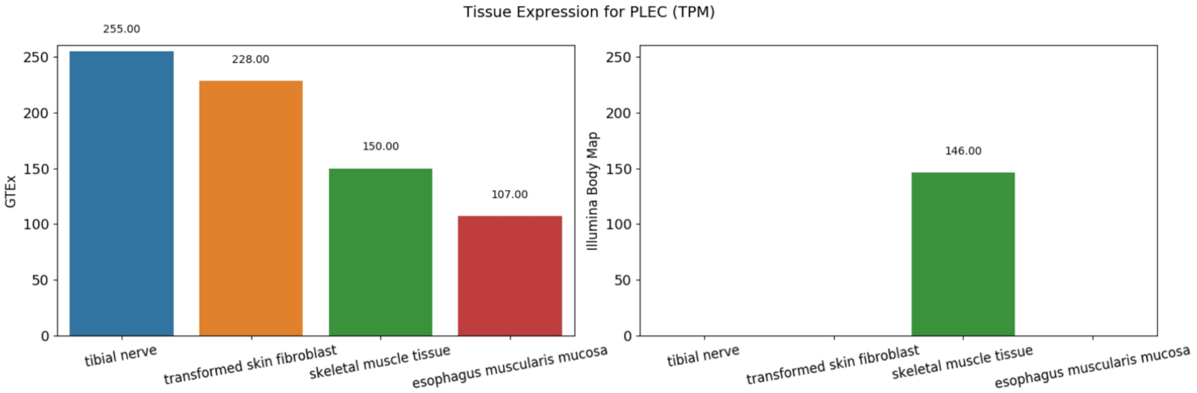
Tissue Expression for PLEC (TPM)
GTEx
Illumina Body Map
0
50
100
150
200
250
255.00
228.00
150.00
107.00
146.00
tibial nerve
transformed skin fibroblast
skeletal muscle tissue
esophagus muscularis mucosa

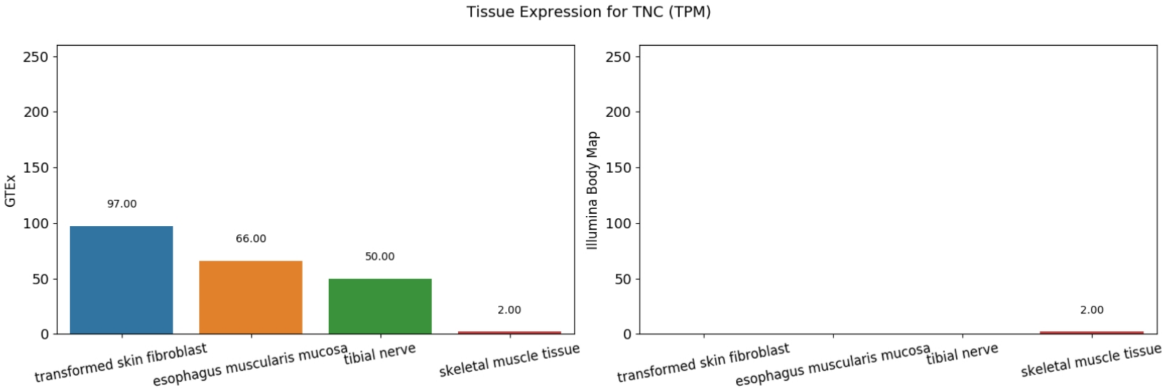
Tissue Expression for TNC (TPM)
GTEx
Illumina Body Map
0
50
100
150
200
250
97.00
66.00
50.00
2.00
2.00
transformed skin fibroblast
esophagus muscularis mucosa
tibial nerve
skeletal muscle tissue

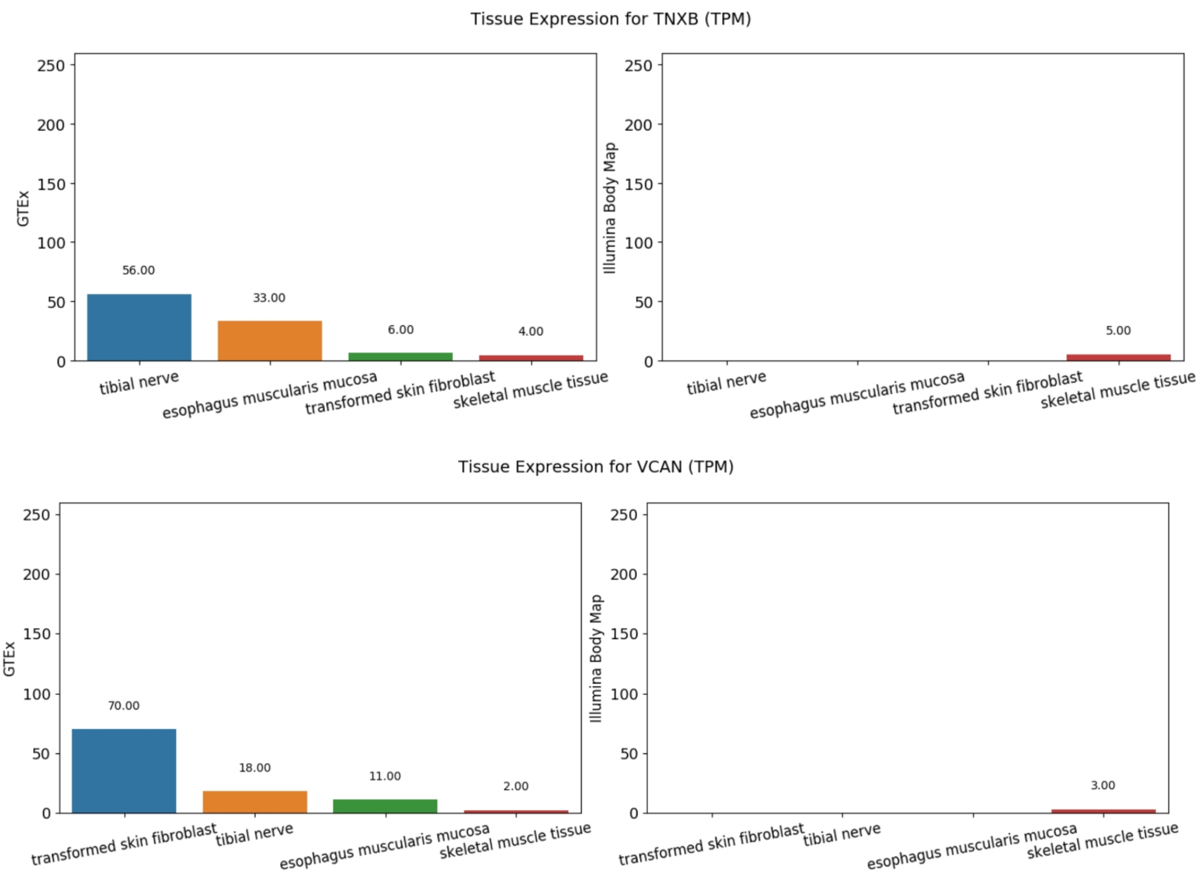


**Suppl. Figure 8.** Tissue-specific expression levels (Transcripts Per Million, TPM) of the genes contained in the largest module within the multilayer communities of the severe group (Methods). Expression levels are reported for GTEx (left panels) and Illumina Body Map (right panels) using EBI Expression Atlas default cutoff (0.5 TPM). Missing bars indicate no data availability (e.g. *COL13A1, LOXL3*). As its expression is below the cutoff for the tissues of interest in both GTEx and Illumina Body Map (Methods), *USH2A* is not reported. Expression level categories based on Expression Atlas: low (0.5 to 10 TPM), medium (11 to 1000 TPM), and high (more than 1000 TPM).

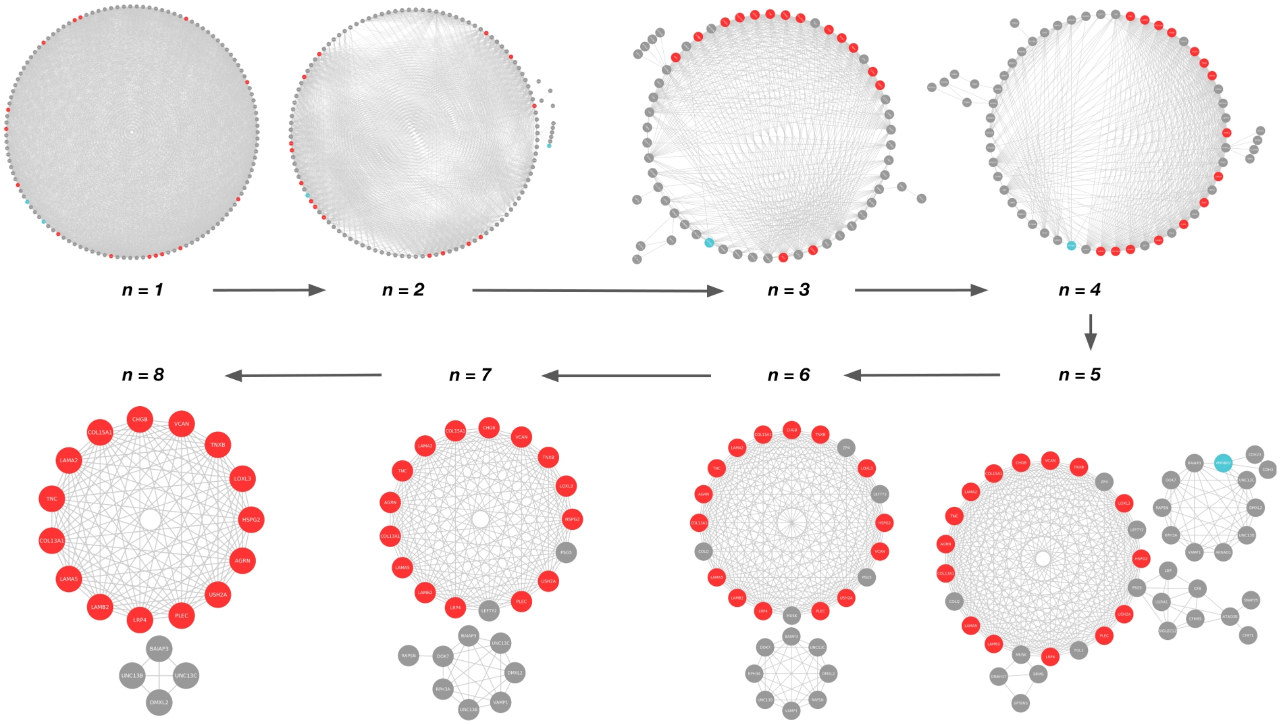


**Suppl. Figure 9.** Presence of *PPFIBP2* and *ACOT2* in the multilayer communities across the range of resolution parameter values (Methods). Genes of the severe-specific module are highlighted in red. *PPFIBP2* (present from n=1 to n=5) and *ACOT2* (present from n=1 to n=2) are depicted in blue.

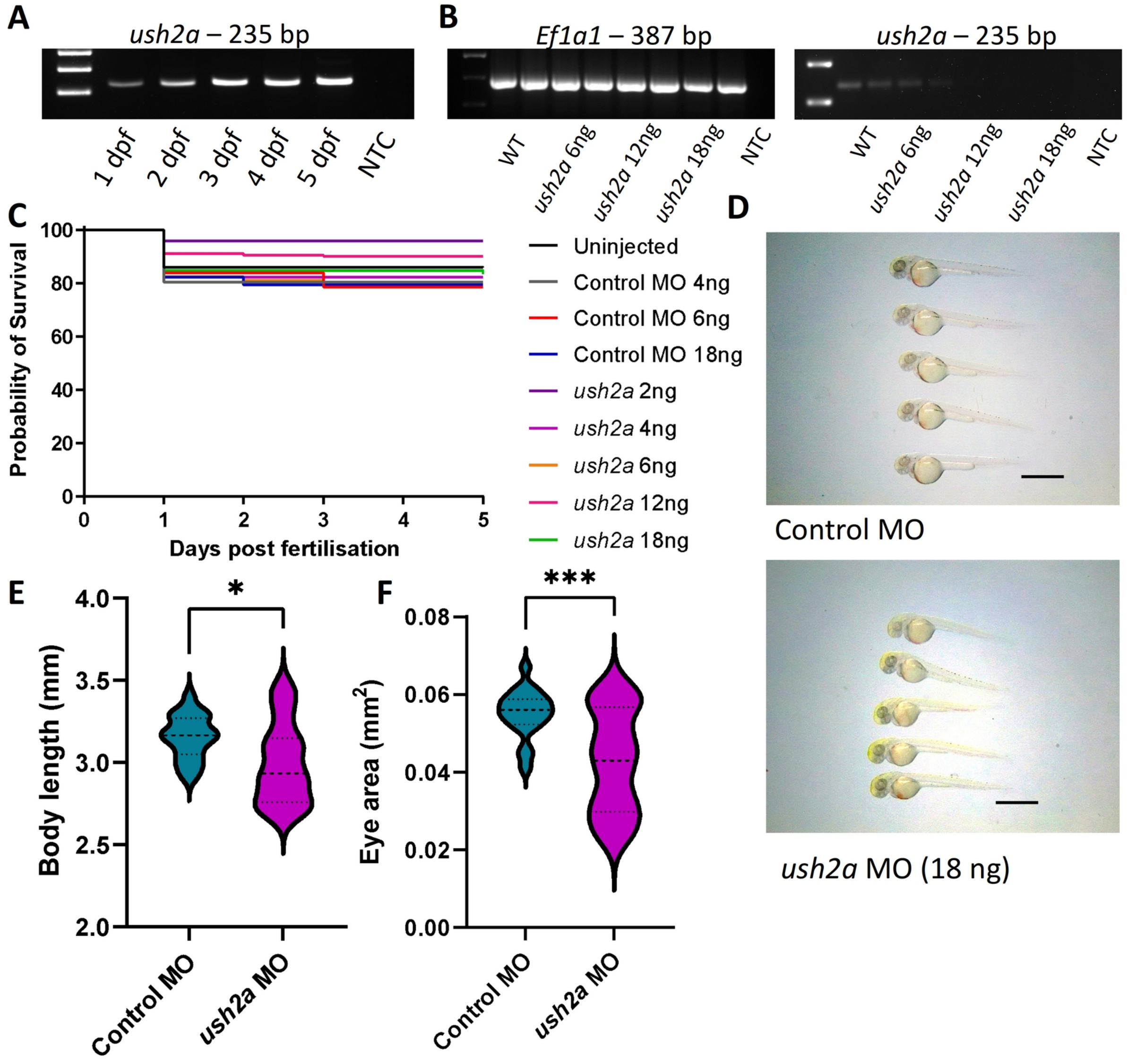


**Suppl. Figure 10.** Survival and phenotype of Ush2a-MO zebrafish A) Reverse-transcriptase (RT)-PCR of wildtype (WT) zebrafish at 1-5 days post fertilization (dpf) showing expression of *ush2a* throughout early development. (B) RT-PCR of control *ush2a*-MO fish at 2 dpf showing consistent expression of eef1a1l1 and a loss of expression of *ush2a* in *ush2a*-MO fish when injected with 6, 12 and 18 ng of MO. NTC = no template control. (C) Survival of WT, control MO and *ush2a*-MO injected zebrafish over 5 dpf. (D) Example light microscope images of control MO and 18 ng *ush2a*-MO-injected zebrafish at 2 dpf. Scale bar = 2mm. (E) Length of 2 dpf control and *ush2a*-MO zebrafish from the tip of the head to the tail. (F) Eye area of control and *ush2a*-MO zebrafish at 2 dpf. Dashed line shows the median, dotted lines show the quartiles, *p < 0.05, ***p < 0.001, unpaired t-test.

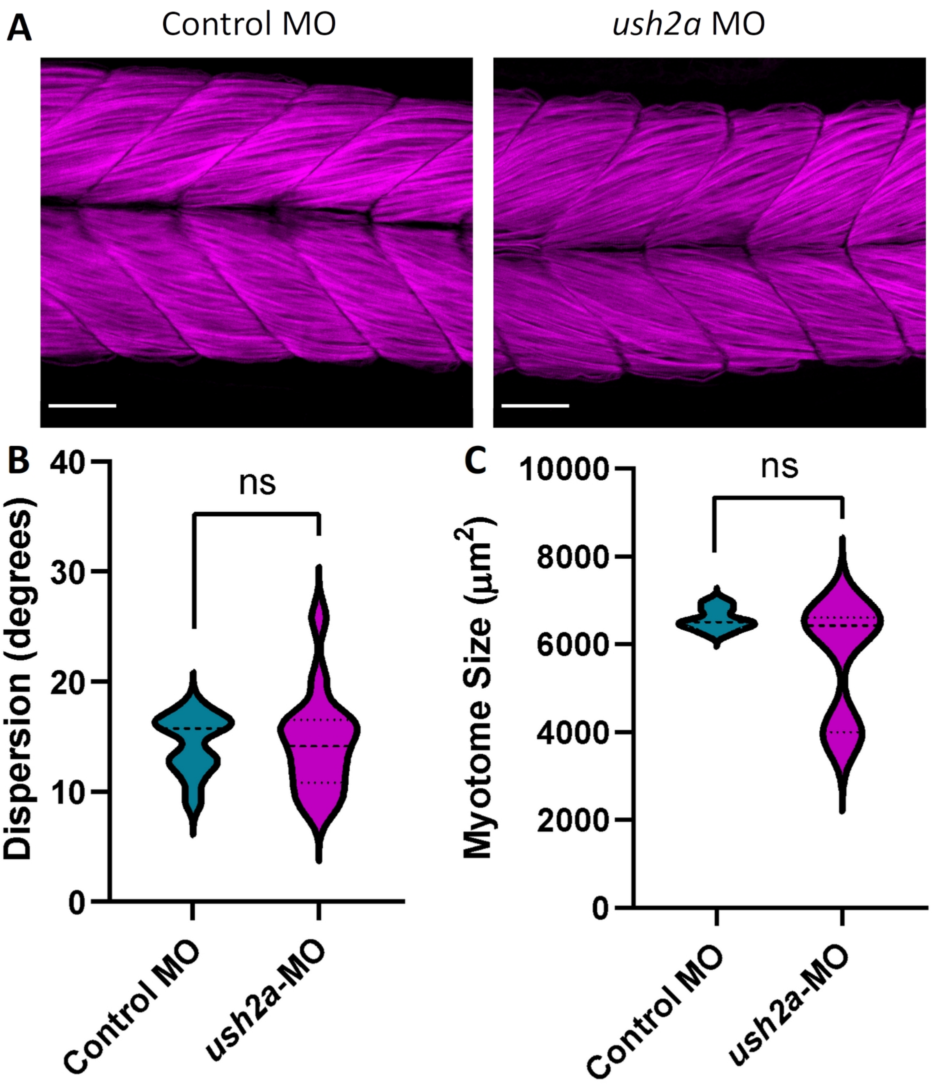


**Suppl. Figure 11.** Muscle morphology in *ush2a*-MO zebrafish. (A) Representative images of phalloidin-stained muscle fibers (detects filamentous-actin) in control and *ush2a* MO 2 dpf fish. Regularly arranged muscle fibers can be observed, with no obvious indications of disorganization, missing fibers or fiber size changes. Scale bar = 50 µm. (B) Dispersion, as a measure of muscle fiber orientation and arrangement, showed no significant differences between the two groups. (C) Myotome size was also similar in control and *ush2a*-MO injected zebrafish. Dashed line shows the median, dotted lines show the quartiles, ns = not significant, nested t-test/unpaired t-test.

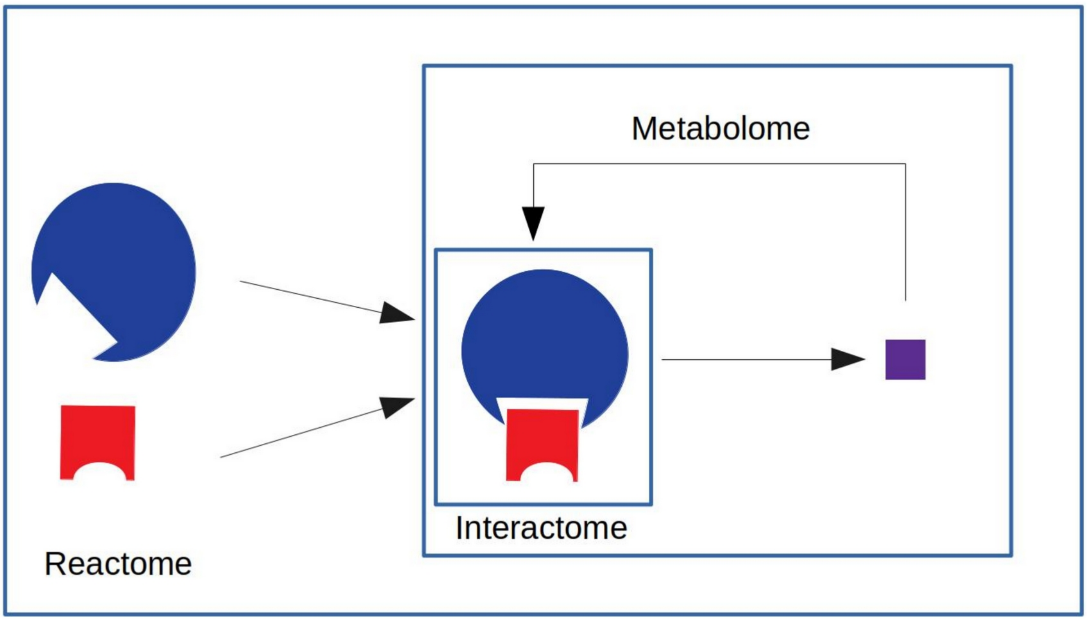


**Suppl. Figure 12.** Distinct layers of biological information covered in the analysis. In this example, enzyme A physically interacts (**interactome)** with enzyme B for the production of a metabolite that is further processed (**metabolome**). Moreover, all these molecules are part of the same pathway (**reactome**).

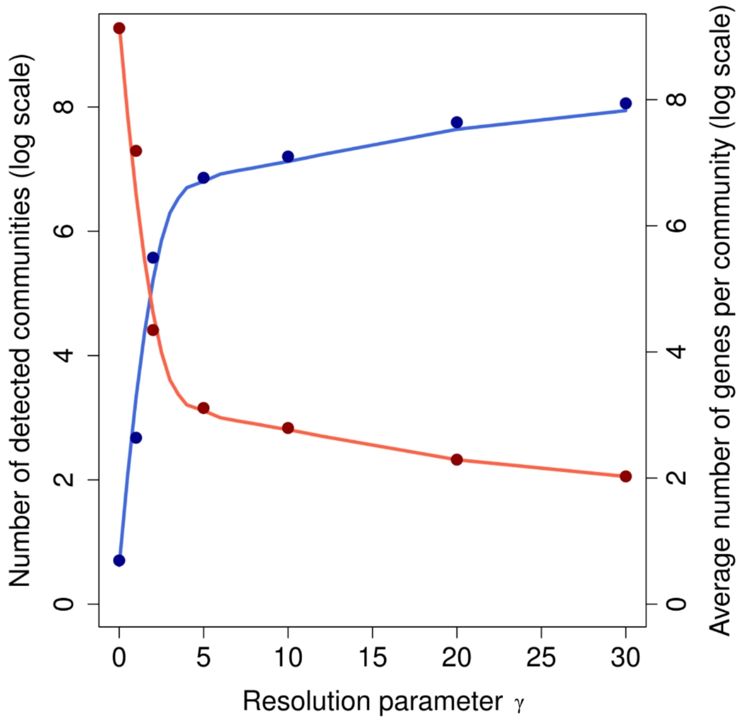


**Suppl. Figure 13.** Variation of number (blue line) and size (red line) of detected multilayer communities as a function of the MolTi resolution parameter γ. Curves are fitted with LOESS (locally estimated scatterplot smoothing) regression with span 0.6. Sampled points along the explored interval are shown for ease of visualization.

### Supplementary Table legends

**Suppl. Table 1.** Clinical characterization of 20 CMS patients with distinct severity levels, namely severe and not-severe (mild and moderate). The annotation of clinical test responses with Human Phenotype Ontology (HPO) (https://hpo.jax.org/) terms has been manually curated. FCV: Forced Vital Capacity, i.e. volume of air that can forcibly be blown out after full inspiration. Y: yes. N: no. NI: no information.

**Suppl. Table 2.** Partially segregating mutations. In the table, mutations segregating at least 50% of one group (i.e. 5 out of 8 severe and 6 out of 10 mild patients) are reported (the mutation categories are described in Supplementary Information).

**Suppl. Table 3**. Genes associated with CNVs and compound heterozygous variants in not-severe and severe phenotypes. Severe-specific genes and known CMS causal genes are reported.

**Suppl. Table 4.** Estimated familiar relatedness between the analyzed patients. Only patients presenting positive relatedness (Methods) are shown.

**Suppl. Table 5.** Functional effect prediction of Ensembl VEP (Method) for the compound heterozygous variants found in the largest module within the multilayer communities of the severe group. Deleterious variants are highlighted in bold.

**Suppl. Table 6.** Functional effect prediction of Ensembl VEP (Method) for the compound heterozygous variants found in Patient 3. Deleterious variants are highlighted in bold.

### Appendix: Supplementary information

#### Functions of CMS-associated genes in the neuromuscular junction

##### *Acetylcholine biosynthesis and release*

Acetylcholine, the main neurotransmitter involved in skeletal muscle contraction, is synthesized in the presynaptic neuron, by the choline acetyltransferase (enzyme encoded by *CHAT* gene), using Acetyl-CoA and choline as substrate in the reaction (Nachmansohn and Machado 1943). Compound heterozygous mutations in this gene were identified by Ohno et al. (K. Ohno et al. 2001) causing CMS in 5 patients.

Solute carriers are critical for this process. Three genes encoding this class of transporters have been previously related to CMS and neuromuscular transmission defects, namely *SLC5A7* (Bauché et al. 2016), *SLC25A* (Chaouch et al. 2014), and *SLC18A3* (O'Grady et al. 2016). *SLC5A7* encodes the membrane choline transporter (Okuda and Haga 2000; Apparsundaram, Ferguson, and Blakely 2001). Acetyl-CoA presence is in part dependent on malate exported from mitochondria, by the action of *SLC25A1* transporter (Kaplan, Mayor, and Wood 1993). Finally, after *CHAT* generates the acetylcholine, this is carried into synaptic vesicles by *SLC18A3* gene product, the VAChT transporter (Eiden et al. 2004).

Another CMS causal gene that might have a detrimental effect at presynaptic level is *PREPL*. This gene encodes a serine oligopeptidase essential for the activation of clathrin associated adaptor protein 1 (AP1), which is needed by VAChTr to fill the synaptic vesicles with acetylcholine (Radhakrishnan et al. 2013). Régal et al. (Régal et al. 2014) described a CMS case caused by a heterozygous deletion. Rabphilin 3a (*RPH3A*) is also involved in vesicle trafficking in the presynaptic element (Guillén et al. 2013; Shirataki et al. 1993) and has recently been described as causative of a specific form of CMS (Ricardo A. Maselli et al. 2018).

Other genes described as causal of CMS related to the vesicle generation and exocytosis are *SNAP25* (Shen et al. 2014), *VAMP1* (Salpietro et al. 2017; Shen et al. 2017), *SYT2* (Herrmann et al. 2014; Whittaker et al. 2015) and *UNC13B* (Andrew G. Engel et al. 2016). *SNAP25* encodes synaptosomal-associated protein 25 (Sørensen et al. 2003), which is a part of the SNARE complex, where also synaptobrevin 1 (*VAMP1*) is allocated (Liu, Sugiura, and Lin 2011). This SNARE complex is key for the $Ca^{2+}$-induced exocytosis of synaptic vesicles, a process in which Synaptotagmin 2 (*SYT2*), the Ca2+ sensor, is also critical (Pang et al. 2006). *UNC13B* encodes a homolog protein to rat Munc13-1. This protein has a calmodulin site and also regulates synaptic vesicles by mediating in the SNARE complex conformation (Ma et al. 2011, 13).

##### *Acetylcholine Receptor clustering*

While acetylcholine is the main neurotransmitter in the neuromuscular junction contraction process, another important molecule, the proteoglycan agrin (*AGRN*), is released by exocytosis from the motor neuron into the synaptic cleft, where it binds the *LRP4* receptor. A special type of myosin, *MYO9*, is known to affect *AGRN* exocytosis upon depletion, causing a characteristic type of CMS (O'Connor et al. 2016; 2018). *AGRN* binding to *LRP4* leads to MuSK (*MUSK*) self-phosphorylation. Activated MuSK recruits Dok-7 (*DOK7*), which in the end stimulates Rapsyn (*RAPSN*) for AChRs (acetylcholine receptors) clustering at the skeletal muscle fiber membrane

(Burden, Yumoto, and Zhang 2013). MuSK, Dok-7 and Rapsyn absence has been previously reported to result in AChR deficiency, poor neuromuscular junction development and causal of some CMS cases (Chevessier et al. 2004; Azulay et al. 1994; Kinji Ohno et al. 2002; Kumar et al. 2018). Interestingly, promoting MuSK activity has been described as capable of preserving neuromuscular synapses in Amyotrophic Lateral Sclerosis mice models (Cantor et al. 2018).

Plectin, encoded by the gene *PLEC*, is essential in the AChR clustering process as it bridges AChRs to the postsynaptic intermediate filament network (IF) via interaction with rapsyn (Mihailovska et al. 2014). Mutations in this gene are also described to cause CMS (Banwell et al. 1999; Selcen et al. 2011). MuSK is also required for the anchoring of endplate acetylcholinesterase (AChE) at the NMJ extracellular matrix (ECM), via a collagen-like peptide encoded by *COLQ* gene (Cartaud et al. 2004). AChE is involved in terminating impulse transmission, by hydrolysis of acetylcholine. Mutations in *COLQ* have been reported as causative for a specific form of CMS (K. Ohno et al. 1998; Donger et al. 1998).

The acetylcholine receptor itself is the main source of CMS-related mutations. In adult individuals, the receptor acts as a cation ligand-gated ion channel formed by 5 homologous subunits, being $\alpha 2\beta\delta\varepsilon$ its stoichiometry. The channel is mainly permeable to $Na^+$ and $K^+$, and to $Ca^{2+}$ in a lesser way. When acetylcholine binds to AChR, the channel opens triggering the membrane depolarization (Brisson and Unwin 1985). All the genes encoding the receptor subunits (*CHRNA, CHRNB, CHRND, CHRNE* and *CHRNG*) have been described as causal for different CMS types (A. G. Engel et al. 1982; Quiram et al. 1999; Brownlow et al. 2001; K. Ohno et al. 1995; Morgan et al. 2006). *CHRNE*, which encodes the ε subunit of the AChR receptor, is causative for ~50% of all reported CMS cases, although frequencies might vary depending on ethnicity (Abicht et al., 1993; Finsterer, 2019). The high prevalence of ε subunit mutations may be the result of partial compensation of its functionality by the embryonic γ (encoded by *CHRNG*), which is substituted after birth given its lower conductance levels. Mutations in other subunits reduce patient survival as no compensation mechanism occurs (Engel et al., 1996). Both Fast-Channel CMS (abnormally short AChR opening time) and Slow-Channel (abnormally long AChR opening time) CMS have been reported for mutations on *CHRNE.*

*SCN4A* gene encodes the α subunit of the voltage-gated sodium channel ($Na_v1.4$), which is key for the generation and propagation of action potentials through the skeletal muscle fiber, which causes $Ca^{2+}$ release and fiber contraction. Many *SCN4A* mutations have been associated with different muscle channelopathies (Wu et al. 2016; Zaharieva et al. 2016; Tsujino et al. 2003), including CMS.

Another process related to AChR clustering is the Endoplasmic Reticulum glycosylation pathway. Normally, mutations in genes that are part of these processes cause congenital disorders of glycosylation (CDG) (Jaeken and Matthijs 2009). However, mutations in some of the pathways components (*DPAGT1*, *ALG2*, *ALG14* , *GFPT1* and *GMPPB*) have been also described as causal of some CMS variants (Belaya et al. 2012; Cossins et al. 2013, 2; Senderek et al. 2011; Belaya et al. 2015).

As for the ECM, collagens are also involved in the receptor clustering process. Collagen XIII (encoded by COL13A1 gene) is known to be a key regulator of NMJ maturation process and AChR clustering (Latvanlehto et al. 2010). Logan et al. (Logan et al. 2015, 19) reported a specific form of CMS being caused by mutations on this gene. Laminins α5 and β2 are also involved molecules in AChR clustering (Rogers and Nishimune 2017). Each one of the different laminins have its own role during NMJ maturation and development, with mutations in LAMA5 (Ricardo A. Maselli et al. 2017) and LAMB2 (R. A. Maselli et al. 2009, 2) being causative of the CMS disease.

**Segregation analyses**

We employed Rbbt (Vázquez et al. 2010) framework to stratify CMS patients based on mutations, aiming to assess whether nonsevere (n=12) and severe (n=8) patients segregate any of the following mutation types (Figure S1):

'**overlapping**' = the mutation overlaps the span of the gene, from first exon to last, including introns
'**mutated_isoform**' = the mutation produces a mutated isoform, i.e. an AA change (on one isoform or just the principal isoform, depending on the options used)
'**splicing**' = the mutation falls within a splicing site, they are deemed to break the protein function
'**affected**' = the mutations affects the encoded protein, by introducing a mutated isoform or a splice site mutation
'**damaged_mutated_isoform**' = the mutation makes a specific protein isoform damaged as predicted by damage or pathogenicity predictions
'**broken**' = the mutation seems to break the protein function, due it introducing a damaging mutation or a splice site mutation
'**TSS**' = the mutation falls within a transcription starting site (1000 bases from TSS)
'**compound**' = the gene has at least two mutations that affect it
'**homozygous**' = the gene is affected by a homozygous mutation
'**missing**' = the genes function may be entirely missing due to a homozygous or a compound mutation possibly affecting both alleles
'**gc19_pc**' **.promCore**' = core promoter of protein coding gene (hg19)
'**gc19_pc.promDomain**' = promoter domain of protein coding gene (hg19)
'**gc19_pc.5utr**' = 5'UTR of a protein coding gene (hg19)
'**gc19_pc.3utr**' = 3'UTR of a protein coding gene (hg19)
'**gc19_pc.ss**' = splicing site of a protein coding gene (hg19)
'**lncrna.promDomain**' = core promoter of a long noncoding RNA with coding potential
'**lncrna.promCore**' = core promoter of a long noncoding RNA with coding potential
'**lncrna.ss**' = splicing site of a long noncoding RNA with coding potential
'**lncrna.ncrna**' = long noncoding RNA
'**smallrna.ncrna**' = small RNA

We define complete segregating mutations that are present in one group and not in the other. No complete segregating mutations were observed (Figure S1), while partial segregation mutations (i.e. present in at least 50% of the patients of one group and not in the other) can be appreciated (Suppl. Table 2).

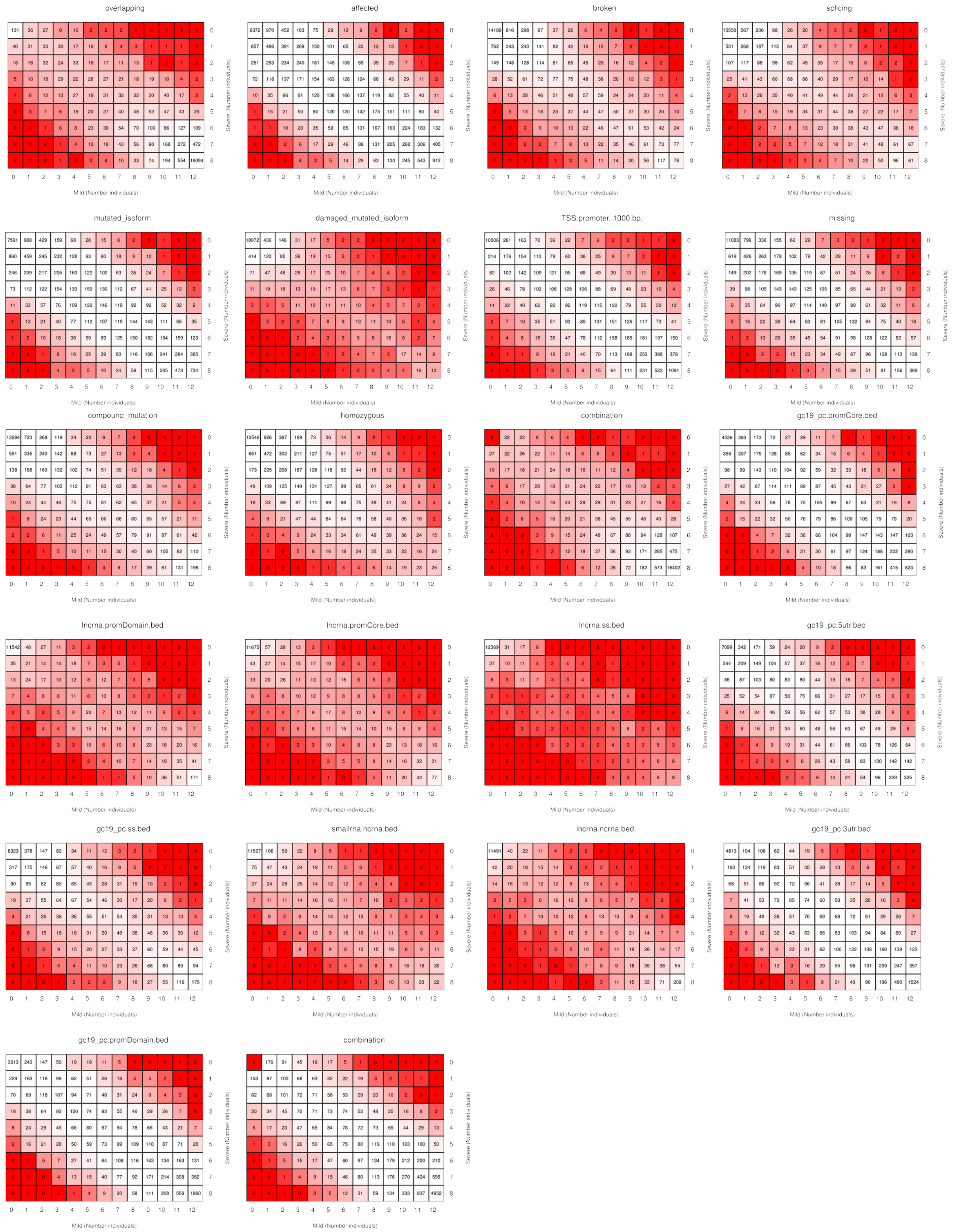


**Figure S1**. Segregation analysis of several mutation types (described in the text). The number of mutations that overlap in the two groups (Nonsevere and Severe) are reported for sets of individuals (0 to 12 for nonsevere individuals, 0 to 8 for severe individuals).

### Multilayer community detection analysis

In this work, we performed a multilayer community detection analysis using MolTi software (Didier, Brun, and Baudot 2015), which adapts the Louvain clustering algorithm with modularity maximization to multilayer networks. The algorithm is parametrized by the resolution parameter $\gamma$: the higher the value of $\gamma$, the smaller the size of the detected multilayer communities.

Given the intrinsic resolution limit of modularity, the reliability of community detection should be assessed *ad hoc* using quality functions that are able to capture the actual community structure of a network (Fortunato and Barthelemy 2007). In this work, we were interested in the identification of communities that robustly express functional relationships among the CMS linked genes (i.e. known CMS causal genes, and severe and nonsevere compound heterozygous variants and CNVs). Accordingly, we sought to determine the largest module of CMS linked genes that are found in the same multilayer communities at any value of resolution within the parameter range in which the community structure is more variable (see Supplementary Figure 12). The adopted procedure is illustrated in Figure S2.

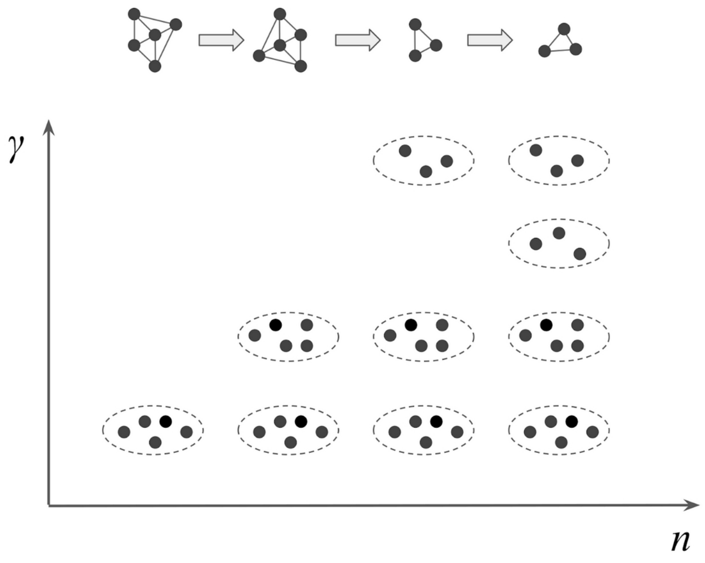


**Figure S2**. Module identification based on detected multilayer communities. Genes that are found in the same community at *n* values of the resolution parameter $\gamma$ are represented as fully connected modules (upper panel). The resolution range considered is $\gamma \in (0,4]$ with intervals of 0.5 (Methods). The module corresponding to the highest *n* contains genes that are systematically found in the same community across the entire range of resolution.

### Supplementary Information References


Abicht, A., Müller, J.S., Lochmüller, H., 1993. Congenital Myasthenic Syndromes Overview, in: Adam, M.P., Mirzaa, G.M., Pagon, R.A., Wallace, S.E., Bean, L.J., Gripp, K.W., Amemiya, A. (Eds.), *GeneReviews®*. University of Washington, Seattle, Seattle (WA).

Apparsundaram, S., S. M. Ferguson, and R. D. Blakely. 2001. 'Molecular Cloning and Characterization of a Murine Hemicholinium-3-Sensitive Choline Transporter'. *Biochemical Society Transactions* 29 (Pt 6): 711–16.

Azulay, J. P., J. Pouget, D. Figarella-Branger, R. Colamarino, J. F. Pellissier, and G. Serratrice. 1994. '[Isolated proximal muscular weakness disclosing myasthenic syndrome]'. *Revue Neurologique* 150 (5): 377–81.

Banwell, B. L., J. Russel, T. Fukudome, X. M. Shen, G. Stilling, and A. G. Engel. 1999. 'Myopathy, Myasthenic Syndrome, and Epidermolysis Bullosa Simplex Due to Plectin Deficiency'. *Journal of Neuropathology and Experimental Neurology* 58 (8): 832–46.

Bauché, Stéphanie, Seana O'Regan, Yoshiteru Azuma, Fanny Laffargue, Grace McMacken, Damien Sternberg, Guy Brochier, et al. 2016. 'Impaired Presynaptic High-Affinity Choline Transporter Causes a Congenital Myasthenic Syndrome with Episodic Apnea'. *American Journal of Human Genetics* 99 (3): 753–61. https://doi.org/10.1016/j.ajhg.2016.06.033.

Belaya, Katsiaryna, Sarah Finlayson, Clarke R. Slater, Judith Cossins, Wei Wei Liu, Susan Maxwell, Simon J. McGowan, et al. 2012. 'Mutations in DPAGT1 Cause a Limb-Girdle Congenital Myasthenic Syndrome with Tubular Aggregates'. *American Journal of Human Genetics* 91 (1): 193–201. https://doi.org/10.1016/j.ajhg.2012.05.022.

Belaya, Katsiaryna, Pedro M. Rodríguez Cruz, Wei Wei Liu, Susan Maxwell, Simon McGowan, Maria E. Farrugia, Richard Petty, et al. 2015. 'Mutations in GMPPB Cause Congenital Myasthenic Syndrome and Bridge Myasthenic Disorders with Dystroglycanopathies'. *Brain: A Journal of Neurology* 138 (Pt 9): 2493–2504. https://doi.org/10.1093/brain/awv185.

Brisson, A., and P. N. Unwin. 1985. 'Quaternary Structure of the Acetylcholine Receptor'. *Nature* 315 (6019): 474–77.

Brownlow, S., R. Webster, R. Croxen, M. Brydson, B. Neville, J. P. Lin, A. Vincent, J. Newsom-Davis, and D. Beeson. 2001. 'Acetylcholine Receptor Delta Subunit Mutations Underlie a Fast-Channel Myasthenic Syndrome and Arthrogryposis Multiplex Congenita'. *The Journal of Clinical Investigation* 108 (1): 125–30. https://doi.org/10.1172/JCI12935.

Burden, Steven J., Norihiro Yumoto, and Wei Zhang. 2013. 'The Role of MuSK in Synapse Formation and Neuromuscular Disease'. *Cold Spring Harbor Perspectives in Biology* 5 (5): a009167. https://doi.org/10.1101/cshperspect.a009167.

Cantor, Sarah, Wei Zhang, Nicolas Delestrée, Leonor Remédio, George Z. Mentis, and Steven J. Burden. 2018. 'Preserving Neuromuscular Synapses in ALS by Stimulating MuSK with a Therapeutic Agonist Antibody'. *ELife* 7. https://doi.org/10.7554/eLife.34375.

Cartaud, Annie, Laure Strochlic, Manuel Guerra, Benoît Blanchard, Monique Lambergeon, Eric Krejci, Jean Cartaud, and Claire Legay. 2004. 'MuSK Is Required for Anchoring Acetylcholinesterase at the Neuromuscular Junction'. *The Journal of Cell Biology* 165 (4): 505–15. https://doi.org/10.1083/jcb.200307164.

Chaouch, Amina, Vito Porcelli, Daniel Cox, Shimon Edvardson, Pasquale Scarcia, Anna De Grassi, Ciro L. Pierri, et al. 2014. 'Mutations in the Mitochondrial Citrate Carrier

SLC25A1 Are Associated with Impaired Neuromuscular Transmission'. *Journal of Neuromuscular Diseases* 1 (1): 75–90. https://doi.org/10.3233/JND-140021.

Chevessier, Frédéric, Brice Faraut, Aymeric Ravel-Chapuis, Pascale Richard, Karen Gaudon, Stéphanie Bauché, Cassandra Prioleau, et al. 2004. 'MUSK, a New Target for Mutations Causing Congenital Myasthenic Syndrome'. *Human Molecular Genetics* 13 (24): 3229–40. https://doi.org/10.1093/hmg/ddh333.

Cossins, Judith, Katsiaryna Belaya, Debbie Hicks, Mustafa A. Salih, Sarah Finlayson, Nicola Carboni, Wei Wei Liu, et al. 2013. 'Congenital Myasthenic Syndromes Due to Mutations in ALG2 and ALG14'. *Brain: A Journal of Neurology* 136 (Pt 3): 944–56. https://doi.org/10.1093/brain/awt010.

Didier, Gilles, Christine Brun, and Anaïs Baudot. 2015. 'Identifying Communities from Multiplex Biological Networks'. *PeerJ* 3: e1525. https://doi.org/10.7717/peerj.1525.

Donger, C., E. Krejci, A. P. Serradell, B. Eymard, S. Bon, S. Nicole, D. Chateau, et al. 1998. 'Mutation in the Human Acetylcholinesterase-Associated Collagen Gene, COLQ, Is Responsible for Congenital Myasthenic Syndrome with End-Plate Acetylcholinesterase Deficiency (Type Ic)'. *American Journal of Human Genetics* 63 (4): 967–75.

Eiden, Lee E., Martin K.-H. Schäfer, Eberhard Weihe, and Burkhard Schütz. 2004. 'The Vesicular Amine Transporter Family (SLC18): Amine/Proton Antiporters Required for Vesicular Accumulation and Regulated Exocytotic Secretion of Monoamines and Acetylcholine'. *Pflugers Archiv: European Journal of Physiology* 447 (5): 636–40. https://doi.org/10.1007/s00424-003-1100-5.

Engel, A. G., E. H. Lambert, D. M. Mulder, C. F. Torres, K. Sahashi, T. E. Bertorini, and J. N. Whitaker. 1982. 'A Newly Recognized Congenital Myasthenic Syndrome Attributed to a Prolonged Open Time of the Acetylcholine-Induced Ion Channel'. *Annals of Neurology* 11 (6): 553–69. https://doi.org/10.1002/ana.410110603.

Engel, A.G., Ohno, K., Bouzat, C., Sine, S.M., Griggs, R.C., 1996. End-plate acetylcholine receptor deficiency due to nonsense mutations in the epsilon subunit. *Annals of Neurology* 40, 810–817. https://doi.org/10.1002/ana.410400521.

Engel, Andrew G., Duygu Selcen, Xin-Ming Shen, Margherita Milone, and C. Michel Harper. 2016. 'Loss of MUNC13-1 Function Causes Microcephaly, Cortical Hyperexcitability, and Fatal Myasthenia'. *Neurology. Genetics* 2 (5): e105. https://doi.org/10.1212/NXG.0000000000000105.

Finsterer, J., 2019. Congenital myasthenic syndromes. *Orphanet Journal of Rare Diseases.* 14, 57. https://doi.org/10.1186/s13023-019-1025-5.

Fortunato, S., and M. Barthelemy. 2007. 'Resolution Limit in Community Detection'. *Proceedings of the National Academy of Sciences* 104 (1): 36–41. https://doi.org/10.1073/pnas.0605965104.

Guillén, Jaime, Cristina Ferrer-Orta, Mònica Buxaderas, Dolores Pérez-Sánchez, Marta Guerrero-Valero, Ginés Luengo-Gil, Joan Pous, et al. 2013. 'Structural Insights into the Ca2+ and PI(4,5)P2 Binding Modes of the C2 Domains of Rabphilin 3A and Synaptotagmin 1'. *Proceedings of the National Academy of Sciences of the United States of America* 110 ( 51): 20503–8. https://doi.org/10.1073/pnas.1316179110.

Herrmann, David N., Rita Horvath, Janet E. Sowden, Michael Gonzalez, Michael Gonzales, Avencia Sanchez-Mejias, Zhuo Guan, et al. 2014. 'Synaptotagmin 2 Mutations Cause an Autosomal-Dominant Form of Lambert-Eaton Myasthenic Syndrome and Nonprogressive Motor Neuropathy'. *American Journal of Human Genetics* 95 (3):

332–39. https://doi.org/10.1016/j.ajhg.2014.08.007.

Jaeken, Jaak, and Gert Matthijs. 2009. 'From Glycosylation to Glycosylation Diseases'. *Biochimica Et Biophysica Acta* 1792 (9): 823. https://doi.org/10.1016/j.bbadis.2009.08.003.

Kaplan, R. S., J. A. Mayor, and D. O. Wood. 1993. 'The Mitochondrial Tricarboxylate Transport Protein. CDNA Cloning, Primary Structure, and Comparison with Other Mitochondrial Transport Proteins'. *The Journal of Biological Chemistry* 268 (18): 13682–90.

Kumar, Ashutosh, Sheila Asghar, Robert Kavanagh, and Matthew P. Wicklund. 2018. 'Unique Presentation of Rapidly Fluctuating Symptoms in a Child with Congenital Myasthenic Syndrome Due to RAPSN Mutation'. *Muscle & Nerve* 58 (4): E23–24. https://doi.org/10.1002/mus.26200.

Latvanlehto, Anne, Michael A. Fox, Raija Sormunen, Hongmin Tu, Tuomo Oikarainen, Anu Koski, Nikolay Naumenko, et al. 2010. 'Muscle-Derived Collagen XIII Regulates Maturation of the Skeletal Neuromuscular Junction'. *The Journal of Neuroscience: The Official Journal of the Society for Neuroscience* 30 (37): 12230–41. https://doi.org/10.1523/JNEUROSCI.5518-09.2010.

Liu, Yun, Yoshie Sugiura, and Weichun Lin. 2011. 'The Role of Synaptobrevin1/VAMP1 in Ca2+-Triggered Neurotransmitter Release at the Mouse Neuromuscular Junction'. *The Journal of Physiology* 589 (Pt 7): 1603–18. https://doi.org/10.1113/jphysiol.2010.201939.

Logan, Clare V., Judith Cossins, Pedro M. Rodríguez Cruz, David A. Parry, Susan Maxwell, Pilar Martínez-Martínez, Joey Riepsaame, et al. 2015. 'Congenital Myasthenic Syndrome Type 19 Is Caused by Mutations in COL13A1, Encoding the Atypical Non-Fibrillar Collagen Type XIII A1 Chain'. *American Journal of Human Genetics* 97 (6): 878–85. https://doi.org/10.1016/j.ajhg.2015.10.017.

Ma, Cong, Wei Li, Yibin Xu, and Josep Rizo. 2011. 'Munc13 Mediates the Transition from the Closed Syntaxin-Munc18 Complex to the SNARE Complex'. *Nature Structural & Molecular Biology* 18 (5): 542–49. https://doi.org/10.1038/nsmb.2047.

Maselli, R. A., J. J. Ng, J. A. Anderson, O. Cagney, J. Arredondo, C. Williams, H. B. Wessel, H. Abdel-Hamid, and R. L. Wollmann. 2009. 'Mutations in LAMB2 Causing a Severe Form of Synaptic Congenital Myasthenic Syndrome'. *Journal of Medical Genetics* 46 (3): 203–8. https://doi.org/10.1136/jmg.2008.063693.

Maselli, Ricardo A., Juan Arredondo, Jessica Vázquez, Jessica X. Chong, University of Washington Center for Mendelian Genomics, Michael J. Bamshad, Deborah A. Nickerson, et al. 2017. 'Presynaptic Congenital Myasthenic Syndrome with a Homozygous Sequence Variant in LAMA5 Combines Myopia, Facial Tics, and Failure of Neuromuscular Transmission'. *American Journal of Medical Genetics. Part A* 173 (8): 2240–45. https://doi.org/10.1002/ajmg.a.38291.

Maselli, Ricardo A., Jessica Vázquez, Leah Schrumpf, Juan Arredondo, Marian Lara, Jonathan B. Strober, Peter Pytel, Robert L. Wollmann, and Michael Ferns. 2018. 'Presynaptic Congenital Myasthenic Syndrome with Altered Synaptic Vesicle Homeostasis Linked to Compound Heterozygous Sequence Variants in RPH3A'. *Molecular Genetics & Genomic Medicine* 6 (3): 434–40. https://doi.org/10.1002/mgg3.370.

Mihailovska, Eva, Marianne Raith, Rocio G. Valencia, Irmgard Fischer, Mumna Al Banchaabouchi, Ruth Herbst, and Gerhard Wiche. 2014. 'Neuromuscular Synapse Integrity Requires Linkage of Acetylcholine Receptors to Postsynaptic Intermediate Filament Networks via Rapsyn-Plectin 1f Complexes'. *Molecular Biology of the Cell* 25 (25): 4130–49. https://doi.org/10.1091/mbc.E14-06-1174.

Morgan, Neil V., Louise A. Brueton, Phillip Cox, Marie T. Greally, John Tolmie, Shanaz Pasha, Irene A. Aligianis, et al. 2006. ‘Mutations in the Embryonal Subunit of the Acetylcholine Receptor (CHRNG) Cause Lethal and Escobar Variants of Multiple Pterygium Syndrome’. *American Journal of Human Genetics* 79 (2): 390–95. https://doi.org/10.1086/506256.

Nachmansohn, D., and A. L. Machado. 1943. ‘THE FORMATION OF ACETYLCHOLINE. A NEW ENZYME: “CHOLINE ACETYLASE”’. *Journal of Neurophysiology* 6 (5): 397–403. https://doi.org/10.1152/jn.1943.6.5.397.

O’Connor, Emily, Ana Töpf, Juliane S. Müller, Daniel Cox, Teresinha Evangelista, Jaume Colomer, Angela Abicht, et al. 2016. ‘Identification of Mutations in the MYO9A Gene in Patients with Congenital Myasthenic Syndrome’. *Brain: A Journal of Neurology* 139 (Pt 8): 2143–53. https://doi.org/10.1093/brain/aww130.

O’Connor, Emily, Vietxuan Phan, Isabell Cordts, George Cairns, Stefan Hettwer, Daniel Cox, Hanns Lochmüller, and Andreas Roos. 2018. ‘MYO9A Deficiency in Motor Neurons Is Associated with Reduced Neuromuscular Agrin Secretion’. *Human Molecular Genetics* 27 (8): 1434–46. https://doi.org/10.1093/hmg/ddy054.

O’Grady, Gina L., Corien Verschuuren, Michaela Yuen, Richard Webster, Manoj Menezes, Johanna M. Fock, Natalie Pride, et al. 2016. ‘Variants in SLC18A3, Vesicular Acetylcholine Transporter, Cause Congenital Myasthenic Syndrome’. *Neurology* 87 (14): 1442–48. https://doi.org/10.1212/WNL.0000000000003179.

Ohno, K., J. Brengman, A. Tsujino, and A. G. Engel. 1998. ‘Human Endplate Acetylcholinesterase Deficiency Caused by Mutations in the Collagen-like Tail Subunit (ColQ) of the Asymmetric Enzyme’. *Proceedings of the National Academy of Sciences of the United States of America* 95 (16): 9654–59.

Ohno, K., D. O. Hutchinson, M. Milone, J. M. Brengman, C. Bouzat, S. M. Sine, and A. G. Engel. 1995. ‘Congenital Myasthenic Syndrome Caused by Prolonged Acetylcholine Receptor Channel Openings Due to a Mutation in the M2 Domain of the Epsilon Subunit’. *Proceedings of the National Academy of Sciences of the United States of America* 92 (3): 758–62.

Ohno, K., A. Tsujino, J. M. Brengman, C. M. Harper, Z. Bajzer, B. Udd, R. Beyring, S. Robb, F. J. Kirkham, and A. G. Engel. 2001. ‘Choline Acetyltransferase Mutations Cause Myasthenic Syndrome Associated with Episodic Apnea in Humans’. *Proceedings of the National Academy of Sciences of the United States of America* 98 (4):2017–22. https://doi.org/10.1073/pnas.98.4.2017.

Ohno, Kinji, Andrew G. Engel, Xin-Ming Shen, Duygu Selcen, Joan Brengman, C. Michel Harper, Akira Tsujino, and Margherita Milone. 2002. ‘Rapsyn Mutations in Humans Cause Endplate Acetylcholine-Receptor Deficiency and Myasthenic Syndrome’. *American Journal of Human Genetics* 70 (4): 875–85. https://doi.org/10.1086/339465.

Okuda, T., and T. Haga. 2000. ‘Functional Characterization of the Human High-Affinity Choline Transporter’. *FEBS Letters* 484 (2): 92–97.

Pang, Zhiping P., Jianyuan Sun, Josep Rizo, Anton Maximov, and Thomas C. Südhof. 2006. ‘Genetic Analysis of Synaptotagmin 2 in Spontaneous and Ca2+-Triggered Neurotransmitter Release’. *The EMBO Journal* 25 (10): 2039–50. https://doi.org/10.1038/sj.emboj.7601103.

Quiram, P. A., K. Ohno, M. Milone, M. C. Patterson, N. J. Pruitt, J. M. Brengman, S. M.

Sine, and A. G. Engel. 1999. 'Mutation Causing Congenital Myasthenia Reveals Acetylcholine Receptor Beta/Delta Subunit Interaction Essential for Assembly'. *The Journal of Clinical Investigation* 104 (10): 1403–10. https://doi.org/10.1172/JCI8179.

Radhakrishnan, Karthikeyan, Jennifer Baltes, John W. M. Creemers, and Peter Schu. 2013. 'Trans-Golgi Network Morphology and Sorting Is Regulated by Prolyl-Oligopeptidase- like Protein PREPL and the AP-1 Complex Subunit M1A'. *Journal of Cell Science* 126 (Pt 5): 1155–63. https://doi.org/10.1242/jcs.116079.

Régal, Luc, Xin-Ming Shen, Duygu Selcen, Chantal Verhille, Sandra Meulemans, John W. M. Creemers, and Andrew G. Engel. 2014. 'PREPL Deficiency with or without Cystinuria Causes a Novel Myasthenic Syndrome'. *Neurology* 82 (14): 1254–60. https://doi.org/10.1212/WNL.0000000000000295.

Rogers, Robert S., and Hiroshi Nishimune. 2017. 'The Role of Laminins in the Organization and Function of Neuromuscular Junctions'. *Matrix Biology: Journal of the International Society for Matrix Biology* 57–58: 86–105. https://doi.org/10.1016/j.matbio.2016.08.008.

Salpietro, Vincenzo, Weichun Lin, Andrea Delle Vedove, Markus Storbeck, Yun Liu, Stephanie Efthymiou, Andreea Manole, et al. 2017. 'Homozygous Mutations in VAMP1 Cause a Presynaptic Congenital Myasthenic Syndrome'. *Annals of Neurology* 81 (4): 597–603. https://doi.org/10.1002/ana.24905.

Selcen, D., V. C. Juel, L. D. Hobson-Webb, E. C. Smith, D. E. Stickler, A. V. Bite, K. Ohno, and A. G. Engel. 2011. 'Myasthenic Syndrome Caused by Plectinopathy'. *Neurology* 76 (4): 327–36. https://doi.org/10.1212/WNL.0b013e31820882bd.

Senderek, Jan, Juliane S. Müller, Marina Dusl, Tim M. Strom, Velina Guergueltcheva, Irmgard Diepolder, Steven H. Laval, et al. 2011. 'Hexosamine Biosynthetic Pathway Mutations Cause Neuromuscular Transmission Defect'. *American Journal of Human Genetics* 88 (2): 162–72. https://doi.org/10.1016/j.ajhg.2011.01.008.

Shen, Xin-Ming, Rosana H. Scola, Paulo J. Lorenzoni, Cláudia S. K. Kay, Lineu C. Werneck, Joan Brengman, Duygu Selcen, and Andrew G. Engel. 2017. 'Novel Synaptobrevin-1 Mutation Causes Fatal Congenital Myasthenic Syndrome'. *Annals of Clinical and Translational Neurology* 4 (2): 130–38. https://doi.org/10.1002/acn3.387.

Shen, Xin-Ming, Duygu Selcen, Joan Brengman, and Andrew G. Engel. 2014. 'Mutant SNAP25B Causes Myasthenia, Cortical Hyperexcitability, Ataxia, and Intellectual Disability'. *Neurology* 83 (24): 2247–55. https://doi.org/10.1212/WNL.0000000000001079.

Shirataki, H., K. Kaibuchi, T. Sakoda, S. Kishida, T. Yamaguchi, K. Wada, M. Miyazaki, and Y. Takai. 1993. 'Rabphilin-3A, a Putative Target Protein for Smg P25A/Rab3A P25 Small GTP-Binding Protein Related to Synaptotagmin'. *Molecular and Cellular Biology* 13 (4): 2061–68.

Sørensen, Jakob B., Gábor Nagy, Frederique Varoqueaux, Ralf B. Nehring, Nils Brose, Michael C. Wilson, and Erwin Neher. 2003. 'Differential Control of the Releasable Vesicle Pools by SNAP-25 Splice Variants and SNAP-23'. *Cell* 114 (1): 75–86.

Tsujino, Akira, Chantal Maertens, Kinji Ohno, Xin-Ming Shen, Taku Fukuda, C. Michael Harper, Stephen C. Cannon, and Andrew G. Engel. 2003. 'Myasthenic Syndrome Caused by Mutation of the SCN4A Sodium Channel'. *Proceedings of the National Academy of Sciences of the United States of America* 100 (12): 7377–82. https://doi.org/10.1073/pnas.1230273100.

Vázquez, Miguel, Rubén Nogales, Pedro Carmona, Alberto Pascual, and Juan Pavón. 2010. 'Rbbt: A Framework for Fast Bioinformatics Development with Ruby'. In *Advances in*

*Bioinformatics*, edited by Miguel P. Rocha, Florentino Fernández Riverola, Hagit Shatkay, and Juan Manuel Corchado, 74:201–8. Berlin, Heidelberg: Springer Berlin Heidelberg. https://doi.org/10.1007/978-3-642-13214-8_26.

Whittaker, Roger G., David N. Herrmann, Boglarka Bansagi, Bashar Awwad Shiekh Hasan, Robert Muni Lofra, Eric L. Logigian, Janet E. Sowden, et al. 2015. 'Electrophysiologic Features of SYT2 Mutations Causing a Treatable Neuromuscular Syndrome'. *Neurology* 85 (22): 1964–71. https://doi.org/10.1212/WNL.0000000000002185.

Wu, Fenfen, Wentao Mi, Yu Fu, Arie Struyk, and Stephen C. Cannon. 2016. 'Mice with an NaV1.4 Sodium Channel Null Allele Have Latent Myasthenia, without Susceptibility to Periodic Paralysis'. *Brain: A Journal of Neurology* 139 (Pt 6): 1688–99. https://doi.org/10.1093/brain/aww070.

Zaharieva, Irina T., Michael G. Thor, Emily C. Oates, Clara van Karnebeek, Glenda Hendson, Eveline Blom, Nanna Witting, et al. 2016. 'Loss-of-Function Mutations in SCN4A Cause Severe Foetal Hypokinesia or "classical" Congenital Myopathy'. *Brain: A Journal of Neurology* 139 (Pt 3): 674–91. https://doi.org/10.1093/brain/awv352.
